# Distinct transcriptional programs define cranial motor neuron subtypes during vertebrate development

**DOI:** 10.64898/2026.07.02.731459

**Authors:** Austin Q. Seroka, Riku Yasutomi, Italia DiChristina, Adam J. Isabella, Cole Trapnell, Cecilia B. Moens

**Author notes:** Equally contributing authors (Austin Q. Seroka, Riku Yasutomi, Italia DiChristina).

## Abstract

Cranial motor neurons (**cMNs)**, form discrete nuclei that control diverse behaviors such as eye movement, feeding, facial expression, and regulation of visceral organ function. However, the developmental programs that drive cMN target choices and functional specialization have not been comprehensively studied in any vertebrate. Here, we present an integrated single-cell RNA-sequencing atlas of zebrafish cMN development and perform extensive validation by HCR *in situ* hybridization. We find that each cranial motor nucleus expresses a distinct transcriptional signature, and in many cases, we identify transcriptional correlates to functional subtypes within individual nuclei. We find that identity often precedes axon targeting, indicating that cranial motor neuron fate is genetically specified early in development. These distinct identities are shaped by the intersection of shared function, rhombomere origin, and developmental time. A cross-species comparison between zebrafish and mouse reveals that these genetic programs are conserved.

## INTRODUCTION

Cranial motor neurons (**cMNs**) arise in discrete nuclei along the vertebrate midbrain and hindbrain (Fig. 1A) and extend axons in specific cranial nerves (CN) to control diverse behaviors. Oculomotor (CN III), trochlear (CN IV), and abducens (CN VI) motor neurons control distinct extraocular muscles, whereas the branchiomotor neurons: trigeminal (CN V), facial (CN VII), glossopharyngeal (CN IX), and vagal (CN X), innervate branchial arch muscles controlling biting, chewing, swallowing and facial expression. Hindbrain visceromotor neurons additionally provide parasympathetic innervation to glands and organs, or innervate sensory organs to suppress responses to sound and movement^1–3^.

**Fig. 1:**
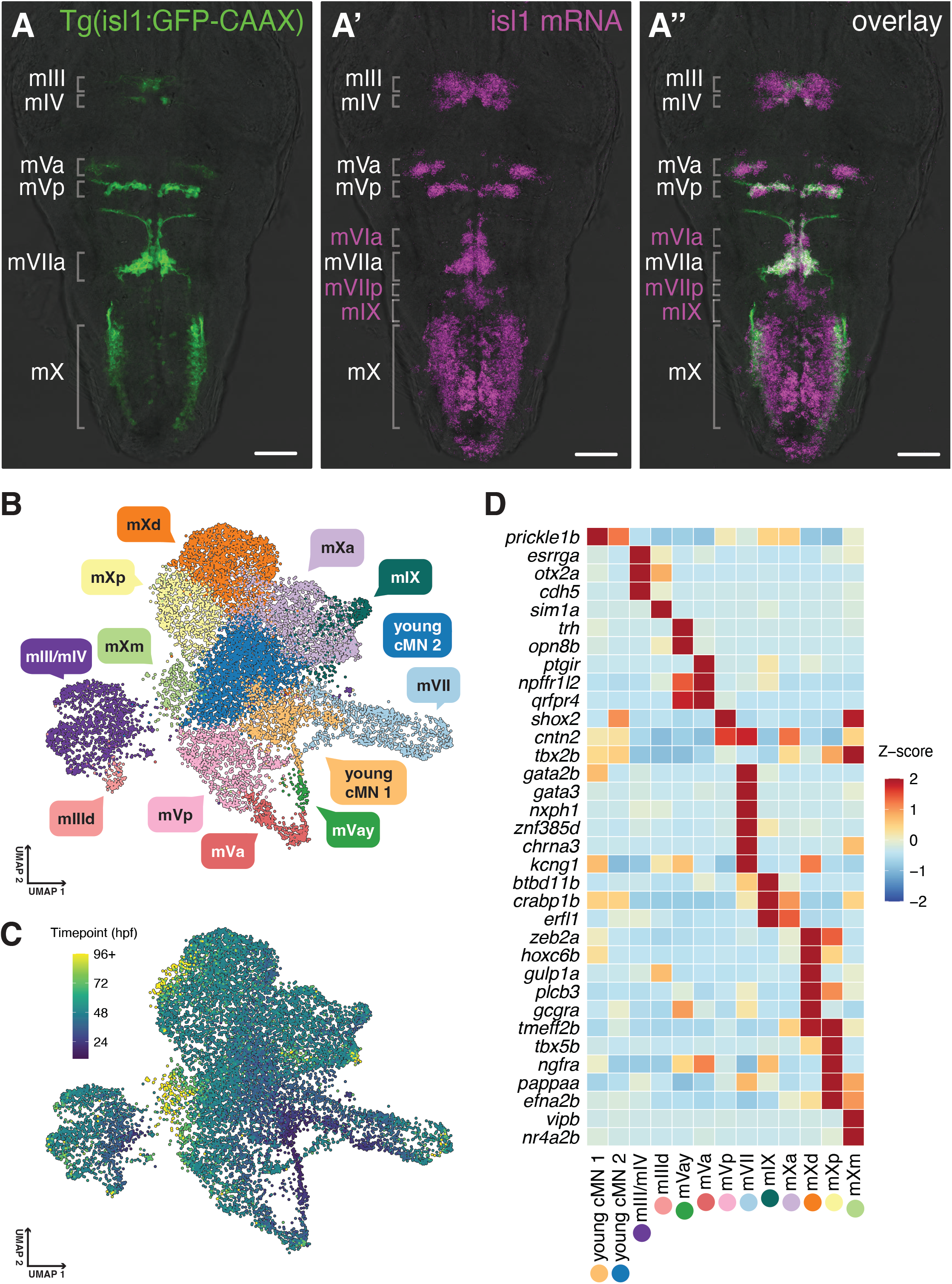
Identification of genes expression differences among cranial motor neurons. **(A-A’’)** Dorsal view of the zebrafish brain at 3 dpf (Max intensity projections; anterior to the top). (A) Tg(*isl1*:GFP-CAAX) (green) labels the mIII, mIV, mVa, mVp, mVIIa and mX motor nuclei. (A’) *isl1a* HCR (magenta) additionally labels the mVI, mVIIp and mIX nuclei, which do not reliably express Tg(*isl1*:GFP:CAAX). (A’’) Overlay. Scale bars 100µm. **(B) UMAP** of the integrated cMN atlas colored and labeled by cMN type determined by HCR. In addition to the nuclei labeled in (A’’), we identified young cMNs, dorsal occulomotor (mIIId), young anterior trigeminal (mVay), anterior (mXa), dorsal (mXd), posterior (mXp), and medial vagus (mXm) identities. **(C)** UMAP colored by sampling timepoint (hpf). **(D)** Heatmap of genes expression in cMNs. Mean log-normalized expression z-scored across cMN types, and clamped to [+2, −2].

cMNs are specifically affected in human congenital cranial dysinnervation disorders in which characteristic defects in eye and facial movements are caused by improper cMN specification, migration and axon guidance^4,5^. cMN and autonomic developmental programs are also related to congenital central hypoventilation syndrome, a clinically severe disruption of respiratory regulation^6,7^. Defining the identities of individual cMN nuclei is essential for uncovering the mechanistic details of motor neuron diversification, and for interpreting how genetic perturbations selectively disrupt the function of individual cranial nerves^8^.

Previous work has uncovered general developmental principles shared across cMNs. The transcription factor Phox2b is required for the formation of branchiomotor and visceromotor neurons^9^. Hox patterning along the anteroposterior axis, specifies hindbrain segment (rhombomere) identity and consequently distinct cMN populations^10,11^. cMNs also use postmitotic identity programs distinct from spinal motor neurons. For example, *Isl1* and *Tbx20* regulate branchiomotor neuron positioning and axon pathfinding^12,13^. However, it remains unclear whether individual cranial motor nuclei can be distinguished by transcriptional identity and whether molecularly distinct subtypes within motor nuclei correlate with functional differences.

Neural identity can be defined by morphology, molecular identity, and function. Single-cell transcriptomics has reshaped how neural cell-types are defined across species. Atlases in mouse and fly link transcriptional identity to developmental origin and anatomical organization within the nervous system^14–20^. Single-nucleus transcriptomics has provided molecular definitions of human cortical cell-types related to disease states^21,22^. In zebrafish, dense developmental sampling has allowed the reconstruction of differentiation trajectories during embryogenesis and larval development^23–26^. These approaches catalogue cell-types and reveal how neural populations arise and diversify over time.

Not all neuronal populations are resolved to the same level of molecular detail. Closely related classes such as motor neurons share core differentiation programs and are difficult to distinguish transcriptionally, despite clear differences in connectivity and function. Single-cell studies of spinal cord diversity have identified molecular distinctions among broad motor neuron classes ^27–30^. Profiling of mouse spinal cord has resolved individual motor pools^28^, whereas zebrafish studies have resolved broad functional classes^31,32^. However, whether closely related motor neuron sub-populations can be distinguished by gene expression remains unresolved and has not been explored in cranial motor neurons.

The zebrafish is well suited to resolve these molecular differences. Optical transparency allows visualization of cMN soma and axonal projections in intact embryos, and genetic tools enable selective labeling of cMNs^33–35^.

Here, we present a scRNA-Seq atlas of zebrafish cMN development from 12 hours to 6 days post-fertilization combining FACS-purified *isl1*^+^ cMNs^36^ generated here with cMNs extracted from two published embryo-scale atlases^25,26^. We validate cluster-specific markers using RNA *in-situ* hybridization,^37^ assessing 51 genes across 4 developmental timepoints in each cMN nucleus. Using these markers we iteratively define clusters corresponding to individual cranial motor nuclei, revealing transcriptional programs that distinguish each nucleus. We further identify transcriptional signatures of motor subtypes within nuclei that correspond to functional groups defined by retrograde labeling. Finally, we perform a cross-species analysis to predict the identities of mouse cMNs, identifying conserved transcriptional features of cranial motor neuron development. Together, this atlas provides a framework for future studies of cranial motor neuron diversification.

## RESULTS

### A single-cell transcriptomic atlas of cranial motor neuron development

Zebrafish cMNs express *isl1a* and transgenes driven by the CREST1 enhancer of *isl1a*^33,38^ (Fig. 1A) (hereafter referred to simply as the *isl1* enhancer). We performed single-cell RNA sequencing (scRNA-seq) of FACS-purified cMNs from Tg(*isl1*:Kaede)^ch_103_^ transgenic hindbrains dissected at 28, 48, 72, and 144 hours post-fertilization (hpf). Purified cells were sequenced using the 10X chromium platform. We selected cMNs by subsetting a *chata+* ^39^); *tbx20+* (cMNs^40–42)^ cluster, composed of 1,114 cells, for further analysis (Fig. S1A).

To increase cMN cell counts, we turned to large-scale whole embryo scRNA-seq datasets. The Trapnell lab single-cell combinatorial indexing (sci-) RNA sequencing atlase spans multiple key developmental timepoints and is composed of millions of cells from control and chemically or genetically perturbed zebrafish embryos^26,43,44^. We first identified clusters corresponding to cMNs (*tbx20+*) and spinal motor neurons (spMNs; *mnx1*+^45^) (Fig. S1B, C) (see methods). Differentially expressed gene (**DEG**) analysis between the *tbx20+* cluster and *mnx1+* cluster identified additional known markers including *phox2bb*^9^ and that distinguish cMNs from spMNs, supporting the cranial identity of the *tbx20+* cluster, composed of 9,695 cells (Fig. S1D). Additionally, we extracted cMNs from the DanioCell 10X chromium atlas^25^ by subsetting the “glia_11” group, processing the cells, and identifying an *isl1+ chata+ tbx20+ mnx1-*cluster, composed of 1,759 cells (Fig. S1E).

The three datasets differ in technical qualities, most notably differences in cell and transcript counts (Fig. S2A, B). The distributions of sampling timepoints were similar between the three datasets, collectively spanning 14 to 144hpf, with the most coverage between 24 and 72hpf (Fig. S2C). To overcome technical differences between the datasets, we applied anchor-based batch correction^46^ to generate an integrated atlas of cMN development composed of 12,568 cells and 9 clusters (Supp Fig. 2D, E; top DEGs listed in Table 1). Some of these clusters were further subclustered to identify additional cMN subtypes (Fig. 1B,C).

**Fig. 2:**
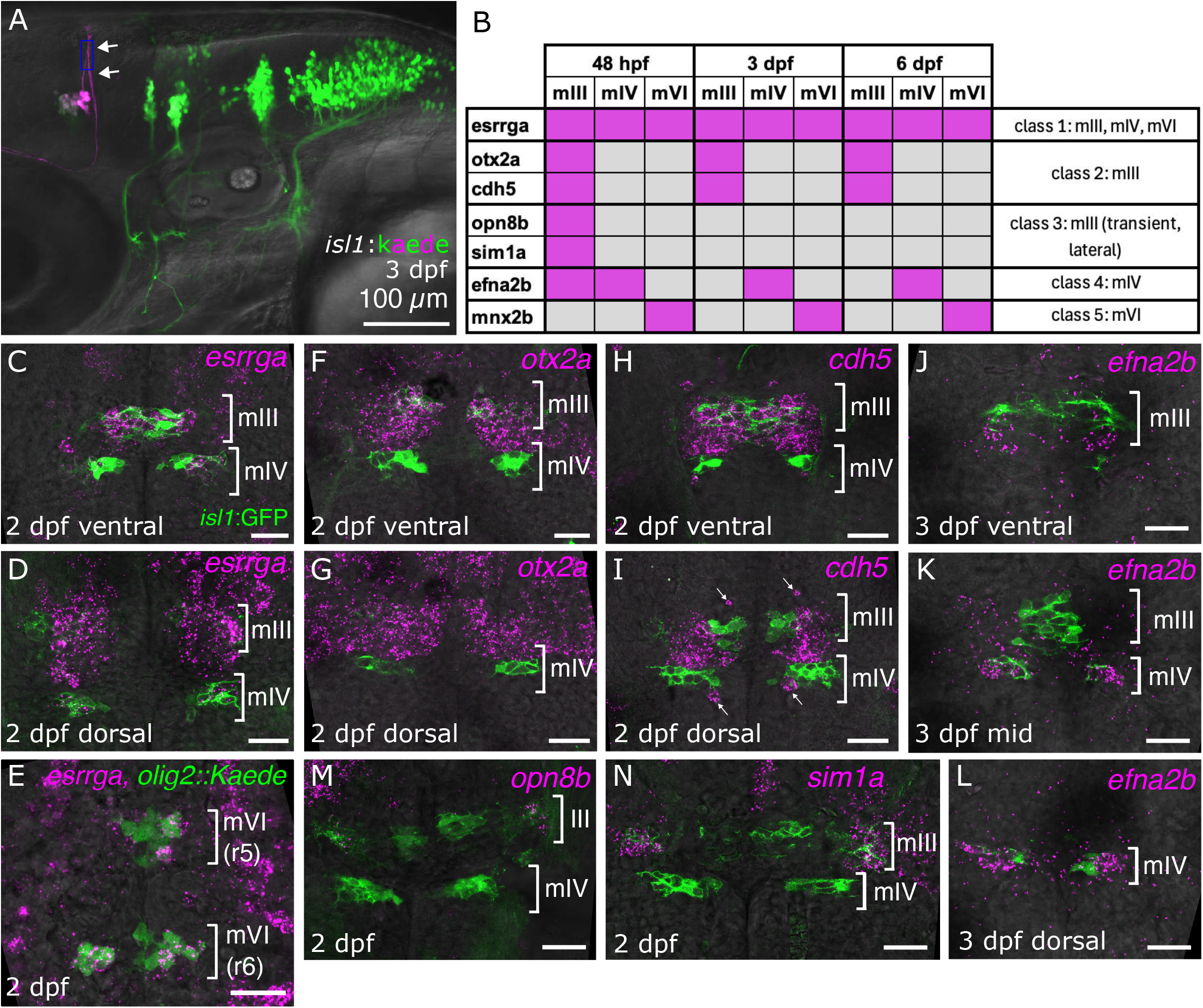
Eye-innervating motor neurons share developmental features distinct from other cMNs. **(A)** Retrograde labeling of trochlear (mIV) cell bodies by photoconversion of their dorsally projecting axon bundles at 3 dpf. Lateral view, anterior to the left. Photoconverted Kaede is magenta and non-photoconverted Kaede is green. Blue box = photoconverted ROI. **(B)** Summary of gene expression in extra-ocular motor neurons as determined by HCR, defining five gene classes. Magenta fill indicates expression; grey fill indicates non-expression.. **(C-N):** HCR in eye-innervating neurons. HCR signal is magenta, Tg(*isl1*:GFP) is green. Single z-sections, ventral view, anterior to the top. “ventral” and “dorsal” sections are ∼12 µm apart. **(C-E**) *esrrga* in mIII (C), mIV (D) and mVI (E) at 2 dpf. *esrrga* in E co-labeled with Tg*(olig2*:Gal4; UAS:Kaede)(green) to define mVI neurons which do not express Tg(*isl1*:GFP). **(F-I)** expression of *otx2a* (F,G) and *cdh5* (H,I) in mIII but not mIV neurons. Arrows in (I) indicate *cdh5* expression in endothelial cells. **(J-L)** expression of *efna2b* in mIV but not mIII neurons at 3dpf. **(M-N)** expression of *opn8b* (M) and *sim1a* (N) in dorso-lateral mIII at 2 dpf. Scale bars = 100µM (A), 20µM (C-N).

Through iterative hybridization chain reaction (**HCR**) screening of cluster and subcluster DEGs, we were able to unambiguously correlate 11 transcriptionally distinct populations with a specific cranial motor nucleus or, in some cases, a functional subdivision of a cranial motor nucleus (Fig. 1B, D and see sections on individual motor nuclei below; top DEGs listed in Table 2). Although this dataset includes cells from genetic and chemical perturbations, perturbed cells were represented in similar proportions across clusters and did not give rise to any novel cluster identities (Fig. S2F).

### Early cranial motor neurons

Two of the cMN states we identify correspond to differentiating cMNs. Cluster 8 is enriched for notch pathway genes *notch1a* and *dlb,* suggesting these are cells that are undergoing or have recently completed neurogenesis (Fig. S1G). Cluster 1 cells are sampled at slightly later timepoints and are less enriched for notch pathway genes compared to cluster 8 (Fig. S2G, H). These cells also lack expression of mature motor neuron transcripts, *elavl4* and *chata,* and have very few DEGs, thus appearing to be differentiating cMNs in a state between cluster 8 and a more mature motor identity (Fig. S2G and Table 1).

### Extra-ocular muscle-innervating motor neurons (CN III, IV and VI)

The oculomotor (III) trochlear (IV) and abducens (VI) nerves innervate the extraocular muscles that move the eye. CN IV and CN VI motor neurons (mIV and mVI neurons) each innervate a single extraocular muscle: the superior oblique (SO) and lateral rectus (LR), respectively. In contrast, CN III motor neurons (mIII) innervate multiple muscles: the superior rectus (SR), inferior rectus (IR), medial rectus (MR), and inferior oblique (IO) muscles^47^. Within mIII, motor neurons are topographically organized in a way that correlates with birth order and function^48,49^. How these anatomical and functional differences between mIII neurons are specified is unknown.

In zebrafish, mIII neurons are born in the ventral midbrain starting at 22 hpf^48^ and begin to express Tg*(isl1*:GFP-CAAX)*^fh474^*, hereafter called *isl1*:GFP) by 26 hpf^33^. mIV neurons are born several hours later, immediately posterior to the midbrain-hindbrain boundary in which they extend their distinctive dorsally crossing axons (Fig. 2A)^33,50^. mVI neurons arise medially in hindbrain rhombomeres 5 and 6^51^. Tg(*isl1*:GFP) expression is more restricted in mIII and mIV neurons than *isl1a* mRNA is and is not detected in mVI neurons at all (Fig. 1A). mVI neurons share developmental and anatomical features with spinal motor neurons: arising from *olig2*+ progenitors^52,53^, expressing *mnx2b*^54^ and having ventral exit points (Fig. S2I)^1^.

Our atlas contains a large, distinct cluster (cluster 4, Fig. S2D) which we identified as the extraocular muscle-innervating neurons by performing HCRs with top DEGs of this cluster at 2 dpf, 3 dpf and 6 dpf (Fig. 2C-L and Fig. S3, showing single horizontal z-sections about 12 microns apart to capture the dorso-ventral dimension of the mIII and mIV nuclei). We identified five classes of genes corresponding to location and timing of expression in these eye-innervating neurons (Fig. 2B).

One of the most specific markers of this cluster, *esrrga,* encoding an orphan nuclear receptor, is detected in all three extraocular muscle-innervating nuclei and not in other cranial motor neurons, although *esrrga* is known to be expressed in other hindbrain neurons^55^ and in spinal motor neurons^31^ (Fig. 2C,D). In both mIII and mIV neurons *esrrga* expression is stronger in outer, *isl1a* mRNA+ neurons than in the core Tg(*isl1*:GFP)+ neurons, particularly at later stages (Fig. S3A-C), possibly reflecting a difference in the developmental state of inner and outer mIII and mIV neurons. Interestingly, *esrrga* is also expressed in mVI neurons, despite their different developmental history (Fig. 2E), suggesting a shared transcriptional code for functionally related motor neurons (see Discussion).

Although mIII and mIV neurons do not cluster separately in our UMAP, HCR validation identified several unique markers of mIII neurons and one unique marker of mIV neurons. Consistent with their origin anterior to the mid-hindbrain boundary, mIII neurons stably express the midbrain determinant *otx2a* while mIV neurons, which develop posterior to this boundary, do not (Fig. 2F,G, Fig. S3D-F). The endothelial cell adhesion marker *cdh5*^56^, is also a specific marker of mIII neurons (Fig. 2H,I, Fig. S3G,H). Additionally, we noted restricted expression of *efna2b*, an ephrin, in mIV neurons but not mIII neurons at 3 and 6 dpf (Fig. 2J-L). *efna2b*-positive cells are widely distributed in our UMAP and *efna2b* is expressed in other cMNs, so without further validation we cannot determine which cells in cluster 4 correspond to mIV neurons.

Subclustering of this large eye-innervating cluster 4 (Fig. S2D) identified eight subclusters (Table 3) of which we characterized only one: a small subcluster (“mIIId” in Fig. 1B; Table 3). HCR validation of *sim1a*, a top DEG in this cluster encoding a transcription factor, showed transient expression in a sub-population of lateral mIII neurons (Fig. 2M). *Opn8b,* an opsin, is also expressed in these neurons (Fig. 2N). Neurons in this position are early-born^48^ and are specifically activated to turn both eyes inward^49^. Since the anatomical tracing that defined these “hunting motor neurons” was done in older larval fish, determining whether the *sim1a* and *opn8b*-expressing neurons correspond to this population will require genetic tools to track their lineage and projections.

A recent preprint used plate-based single-cell RNA-Seq and bulk RNA-Seq of purified extraocular (mIII and mIV) neurons to identify transcriptional programs that might drive their distinct functions in eye movement^57^. Their approach identified many of the same markers that we found as DEGs in cluster 4, as well as an equivalent “mIIId” sub-population (*sim1a^+^, foxp4^+^, pdzrn3b^+^, znf536^+^*), consistent with a distinct transcriptional signature for early-born MR/IR-innervating mIII neurons. They also identified a distinct mIV cluster, the top DEGs for which (*mafba* and *inab*) were not detected as DEGs in our extraocular motor neuron cluster even upon subclustering, possibly because of the shallower sequencing depth in our cells compared to their plate-based approach.

### Trigeminal motor neurons (CN V)

In humans, trigeminal motor (mV) neurons primarily innervate the large muscles derived from the first pharyngeal arch involved in biting and chewing. In fish, trigeminal motor neurons likewise innervate first pharyngeal arch muscles that control jaw movements for feeding and respiration (Fig. S4A,B). A dorsal branch innervates the levator arcus palatini (LAP) and dilator operculi (DO) muscles which expand the buccal cavity, and a mandibular branch passes around the back of the eye and innervates the adductor mandibulae (AM) muscle which closes the mouth. The mandibular branch then continues ventrally to innervate two intermandibularis muscles (IM) which function to raise the floor of the mouth^33,58,59^.

During vertebrate development, mV neurons arise in two bilateral clusters in hindbrain rhombomere 2 (r2; mVa) and rhombomere 3 (r3; mVp) (Fig. 3A)^60^. In zebrafish, mVa becomes Tg(*isl1*:GFP)^+^ by 21 hpf, while mVp arises by 28 hpf^33,61^. Previous retrograde labeling by Di-I injection showed that the mVa and mVp nuclei have distinct muscle targets^33^. We photoconverted Tg(*isl1*:Kaede) in the peripheral branches of the trigeminal motor nerve at their muscle targets at 3 dpf and confirmed these findings: mVa neurons in r2 innervate AM while mVp neurons in r3 innervate LAP/DO dorsally and IM ventrally (Fig. 3B-E). Furthermore, photoconversion of LAP/DO branches predominantly backfilled neurons in a ventro-lateral subdivision of mVp, suggesting the presence of further topographic organization of trigeminal neurons within r3 (Fig. 3C’).

**Fig. 3:**
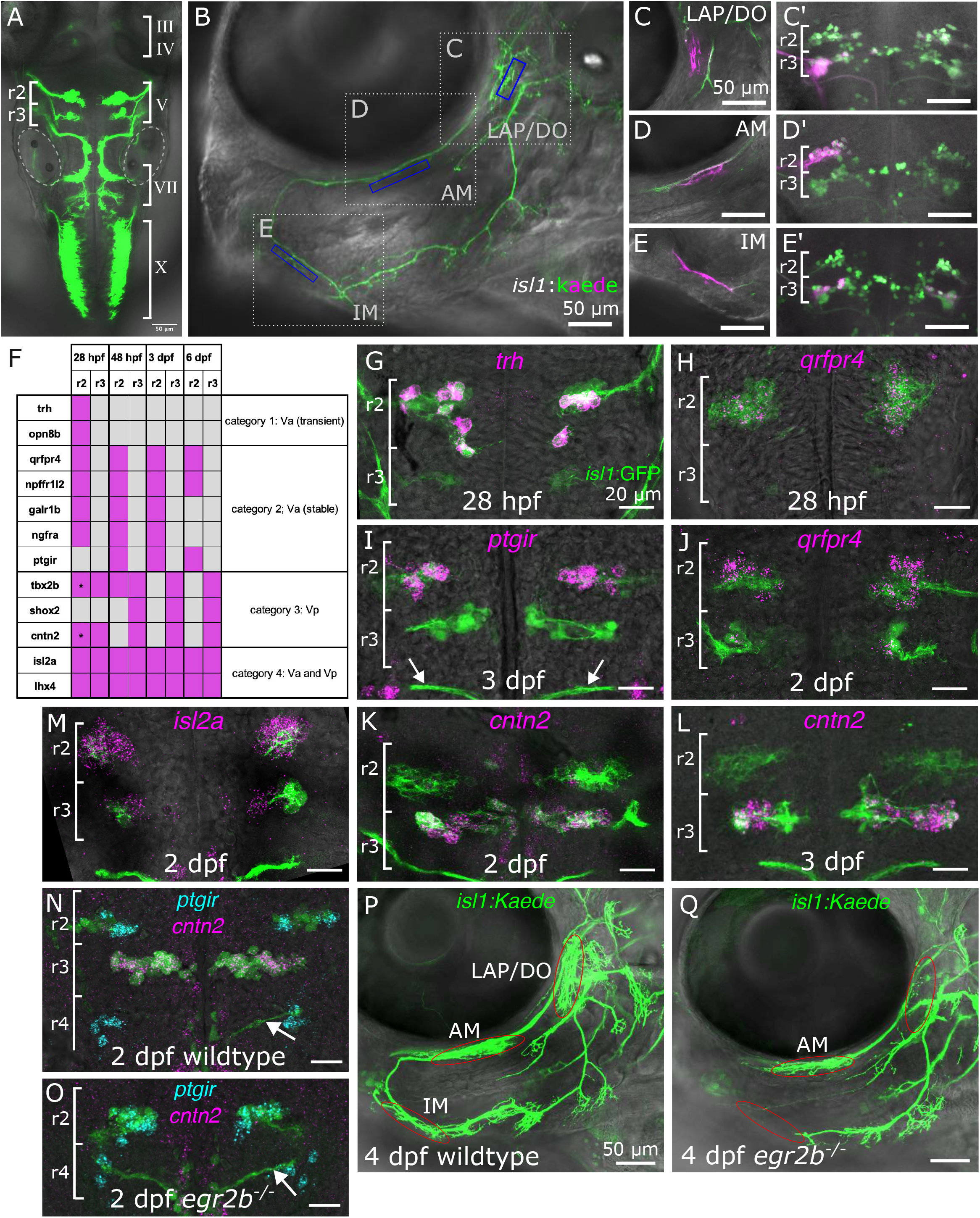
Trigeminal motor neurons have distinct transcriptional identities and muscle targets that depend on early hindbrain rhombomere identity. **(A)** Dorsal view of the cranial motor nuclei in a 2 dpf Tg(*isl1*:GFP)) embryo showing the trigeminal clusters in r2 (mVa) and r3 (mVp). Dotted circles: otic vesicles. **(B)** Ventro-lateral view of a 4 dpf Tg(*isl1*:Kaede) embryo (anterior to the left), showing mV muscle targeting branches. Dotted boxes mark regions imaged in C-E; blue boxes mark ROIs for photoconversion. LAP: levator arcus palatini; DO: dilator operculi; AM: adductor mandibulae; IM: intermandibularis. **(C-E)** Retrograde labeling from mV branches,4 dpf. Photoconverted Kaede is magenta and unconverted Kaede is green. C-E, lateral view of peripheral branches immediately after photoconversion; C’-E’, dorsal view of labeled central cell bodies ∼1 hour later (C-C’). Photoconversion of the LAP/DO branch (C) labels a ventro-lateral subdivision of the ipsilateral mVp (C’). (D-D’) Photoconversion of the AM branch (D) labels ipsilateral mVa (r2) (D’). (E-E’) Photoconversion of the IM branch (E) labels ipsilateral mVp (r3)(E’). **(F)** Summary of gene expression categories in mV neurons. Magenta fill indicates expression. Asterisk: expression in a subset of medial mVa neurons. **(G-M)** HCR validation. Ventral views with anterior to the top. Max projections to capture the ∼30 µm dorso-ventral depth of the mV nuclei. Green: Tg(*isl1*:GFP)); magenta: HCR signal. White arrows indicate the facial (CN VII) motor nerve exiting in r4) **(N-O)** Double HCR for *ptgir* (cyan; mVa) and *cntn2* (magenta; mVp), at 2 dpf. (N) Wild-type embryo; (O) *egr2b^fh227^* homozygous mutant embryo, showing a single *ptgir^+^* mV nucleus. White arrow: CN VII nerve exiting in r4. **(P-Q)** Ventro-lateral views of 4dpf wildtype and *egr2b^fh227^* mutant Tg(*isl1*:Kaede) embryos. Red ovals= muscles innervated by mV neurons (LAP/DO, AM, IM). (P) Wild-type, with all targets innervated. (Q) *egr2b^fh227^* mutant, in which LAP/DO and IM lack mV innervation and only AM remains innervated. Scale bars = 50µm (A-E, P,Q); 20µm (G-O).

Through HCR validation we discovered that mVa and mVp neurons are starkly different in their gene expression profiles (summarized in Fig. 3F). We identified a cluster in our atlas that corresponds specifically to mVa neurons (Fig. 1B). Furthermore, subclustering of cluster 8 (Fig. S2D) distinguished a small subcluster of young (<24 hpf) mVa neurons, mVay (Table 4). Top DEGs of mVay, including *trh* (encoding a pro-hormone) and *opn8b* (an opsin), are detected in mVa neurons at 28 hpf but not at later stages (Fig. 3G, Fig. S4C). mVa markers such as *qrfpr4, npffr1l2,* and *galr1b*, all encoding receptors for neuropeptides, and *ptgir*, a prostaglandin receptor, are specifically and stably expressed in mVa neurons until at least 6 dpf. (Fig. 3H-J and data not shown). We conclude that mVay neurons are at an early stage of their development corresponding to axon outgrowth, while mVa neurons have established stable innervation of the AM muscle.

We identified mVp neurons in a separate cluster (cluster mVp, Fig. 1B). We validated *shox2* and *tbx2b*, both encoding transcription factors, and *cntn2*, encoding a cell adhesion molecule, as expressed in mVp but not mVa neurons (Fig. 3K,L; Fig. S4D-H). *Shox2* is also expressed in a subset of mVII neurons (not shown), suggesting a mixed mVp/mVII identity for this cluster that will be explored in future work. *Tbx2b* and *cntn2* are expressed in other cranial motor nuclei and thus were not identified as DEGs for this cluster (Fig. 1D). Interestingly, within mVp, *tbx2b* expression is restricted to the ventro-lateral cluster that innervates dorsal LAP and DO muscles, suggesting that the topographic relationship of mVp neurons to their specific muscle targets may result from a transcriptional program involving *tbx2b* (Fig. S4G,H).

Pan-cMN genes like *lhx4* and *isl2a* were detected in both mVa and mVp neurons (Fig. 3M). However, no genes were identified that were expressed in both mV nuclei, but not in other cMN populations.

Thus, mVa neurons have a unique transcriptional signature that stably distinguishes them from mVp and all other cMN subtypes, while mVp identity is less uniquely defined. We asked whether these mVa and mVp identities depend on upstream determinants of rhombomere identity. The transcription factor *egr2b* is required for the development of r3 and r5 in mouse and zebrafish, however its role in cranial innervation is unclear^62–64^. At 48 hpf, when wildtype embryos express *ptgir* in mVa and *cntn2* in mVp (Fig. 3N), *egr2b^fh227^* mutants have a single *ptgir*-expressing mV nucleus, consistent with mVa (r2) identity (Fig. 3O; *egr2b^fh227^* is a nonsense allele^65^). Interestingly, none of the muscles normally targeted by mVp (LAP, DO and IM) are innervated in *egr2b^fh227^* mutants, while AM, the muscle normally innervated by mVa neurons, is innervated (Fig. 3P,Q). Thus, mV nuclei are strictly limited in terms of the muscles they can innervate and cannot compensate for each other. The molecular differences between mVa and mVp nuclei are specified early in hindbrain development and correlate with this restricted muscle targeting.

### Facial Motor Neurons (CN VII)

The efferent component of the facial nerve (CN VII) includes the facial branchiomotor neurons (FBMNs) and the octavolateral efferent neurons (OENs) (Fig. 4A,B). In humans, FBMNs innervate the muscles of facial expression, while in fish they innervate the muscles that expand and contract the buccal cavity during suction feeding and respiration^58^ (Fig. 4B white arrows). In mammals, the olivocochlear neurons (OCNs; putatively homologous to OENs) innervate sensory hair cells and spiral ganglia neurons of the ear to protect it from sound-induced damage^66,67^, while in fish, OENs innervate the sensory hair cells of the lateral line and inner ear to suppress reafferent flow signals generated during swimming and to protect vestibular hair cells from overstimulation^68–71^ (Fig. 4B blue arrows). In zebrafish, both FBMNs and OENs are born in hindbrain rhombomere 4 (r4) from which their axons mostly extend, but their cell bodies migrate caudally within the hindbrain from r4 to r6 and r7^34,61^ (Fig. 4A). The first neurons to leave r4 migrate to r7 while later-migrating neurons settle in a large motor nucleus in r6^69,72^. Within the r6 nucleus, a coarse topographic relationship has been described between the dorso-ventral and medio-lateral positions of FBMNs and the muscles they innervate^72^. The OENs that innervate the anterior lateral line neuromasts also lie in r6 and r7, while OENs that innervate posterior lateral line lie primarily in r7^63^. No molecular markers have been identified that distinguish these two types of OENs^70,73^.

**Fig. 4:**
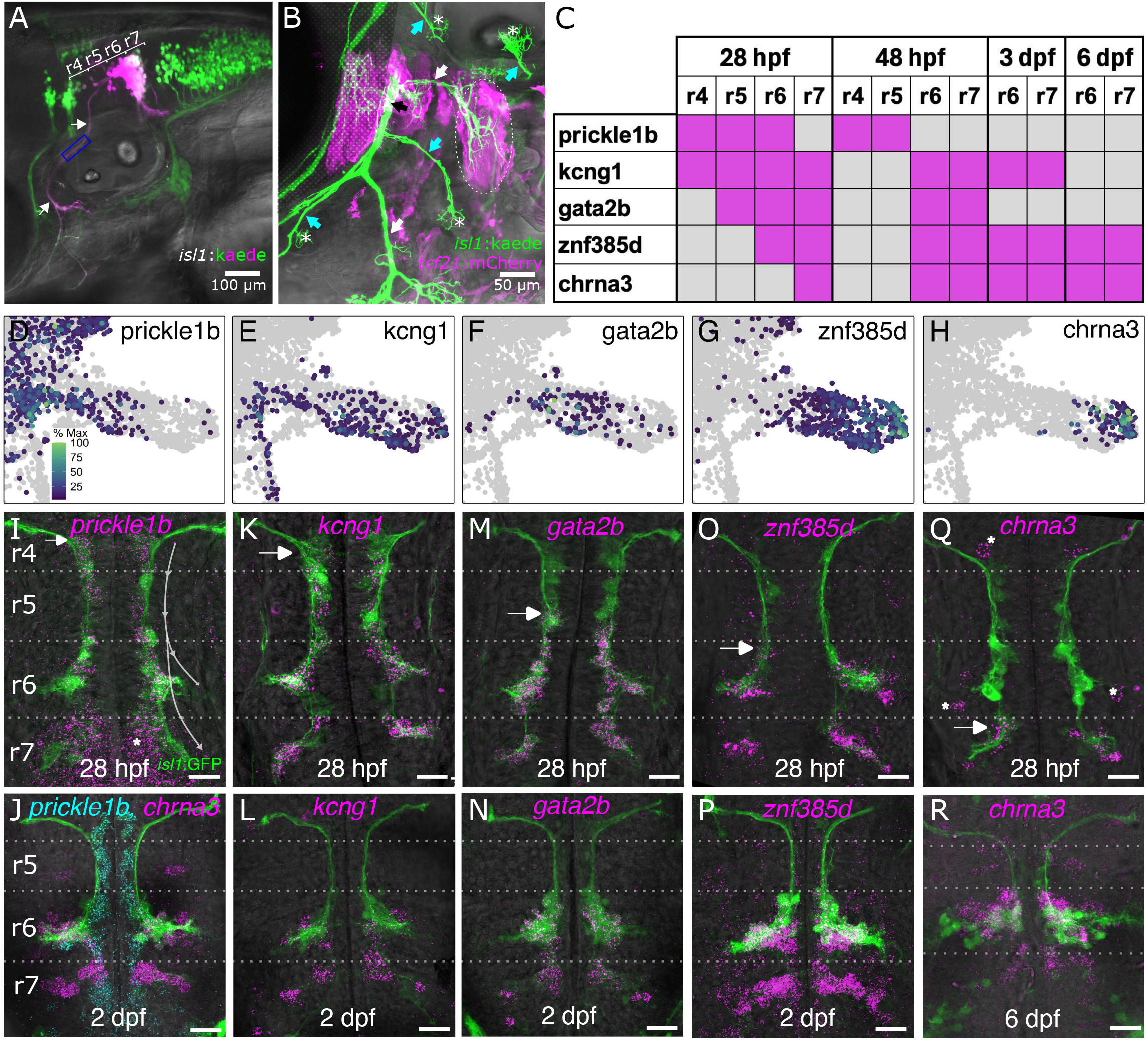
A sequential gene expression program tracks the migration of mVII neurons (A) Lateral view, 3dpf Tg(*isl1*:Kaede) embryo after photoconversion of CN VII (arrows) at ROI indicated (blue box). Photoconverted Kaede (magenta) labels mVII neurons in r6 and r7. Maximum-intensity projection. (B) Lateral view, peripheral branches of mVII neurons at 4 dpf. Neurons express Tg(*isl1*:Kaede) (green) and muscles express TgBAC(*tcf21*:mCherry)(magenta). The mV nerve and its muscle targets are greyed out. Black arrow: main CN VII motor nerve; white arrows: muscle-innervating branches; blue arrows: neuromast-innervating branches; Asterisks: neuromasts. Dotted circle: adductor and levator operculi muscles. (C) Summary of marker expression in mVII neurons. Magenta fill indicates expression. **(D-H)** UMAP of the mVII trajectory (see Fig. 1B) and **(I-R)** HCR validation of genes noted. HCRs are max projections of 20-30 µm z-stacks, ventral views with anterior to the top. Grey arrows in (I) indicate path of mVII neuron migration from rhombomere (r)4 to where they settle in r6 and r7. White arrows in I-R: location of expression onset; dashed lines: rhombomere boundaries; asterisks: non-motor neuron expression. *prickle1b-*expressing cells are at the base of the UMAP trajectory (D) and along the entire migration route but not r6 and r7 (I,J); *kcng1* cells are throughout the trajectory (E) and the entire migration route including r6 and r7 (K,L); *gata2b* cells are mid-trajectory (F) and in r5, r6 and r7 (M,N); *znf385d* cells are mid-to-distal trajectory (G) and in r6 and r7 (O,P); and *chrna3* cells are at the distal tip (H) and initially only in r7 (Q)). Color scale in D-H: percent of the maximum expression level observed for a given gene. Scale bars in I-R: 20 µm.

Herein, we consider OENs and FBMNs together as “mVII neurons”. Future work will describe differences between these populations and their relationship to differences between the OCNs and FBMNs of mammals^74^. We identified a distinct cluster within our atlas corresponding to this population (“mVII” in Fig. 1B). This cluster forms a developmental trajectory in the UMAP embedding with the youngest cells positioned adjacent to the young cMN 1 cluster, and older cells extending towards more mature cell states (Fig. 1C). We identified genes that were expressed at different positions along this UMAP trajectory and validated them by HCR (Fig. 4D-R, summarized in Fig. 4C). We discovered that the UMAP trajectory corresponds closely to phases of mVII neuron migration.

mVII neuron migration requires components of the Planar Cell Polarity pathway^75–78^. Amongst these, *prickle1b* and *nhsl1b* are expressed specifically within migrating mVII neurons and are required cell-autonomously for their migration^79,80^. *prickle1b* and *nhsl1b* are expressed in the youngest cells of the mVII trajectory (Fig. 4D, and data not shown), and we validated *prickle1b* expression by HCR in mVII neurons from the onset of their migration in r4 (Fig. 4I, white arrow). *Prickle1b-*expressing mVII neurons continue to be generated in and migrate from r4 until 2 dpf but *prickle1b* expression is down-regulated after neurons settle in r6 and r7 (Fig. 4J, cyan). *Kcng1,* a K^+^ voltage-gated channel, is also expressed from the onset of mVII migration in r4 at 1 dpf (Fig. 4K, L) but, unlike *prickle1b*, it persists in migrated neurons in r6 and r7 and is turned off in later-migrating neurons at 2 dpf (Fig. 4L). *Gata2b*, a transcription factor that has been implicated in OCN development in mice^81^, is detected further along the mVII trajectory in the UMAP (Fig. 4F), corresponding with onset of expression in mVII neurons only after they have reached r5 (Fig. 4M, white arrow). Like *kcng1*, *gata2b* continues to be expressed in mVII neurons that have settled in r6 and r7 at 2 dpf and is not detected in late-migrating mVII neurons (Fig. 4N). *Znf385d*, encoding one of a large family of zinc finger putative transcription factors, is detected still further along the UMAP trajectory (Fig. 4G) corresponding to onset of expression in mVII neurons only after they have reached r6 (Fig. 4O, white arrow). Finally, chrna3, a nicotinic acetylcholine receptor known to be expressed in OENs^82^, is detected in cells at the very end of the mVII trajectory in the UMAP (Fig. 4H), corresponding to mVII neurons that initiate expression only after they reach their terminal positions in r6 or r7 (Fig. 4Q, white arrow; Fig. 4J (magenta)). *Znf385d* and *chrna3* are terminal markers of mVII neurons, continuing to be expressed at least until our final 6 dpf timepoint (Fig. 4R and data not shown).

Thus, mVII neurons change transcriptionally along the migration path as they mature, in a way that can be predicted by the position of cells in the developmental trajectory of the UMAP. Future work will determine whether these transcriptional changes depend on neuron position or cell-intrinsic developmental timing.

### Glossopharyngeal and Vagus motor neurons: (CN IX and X)

The vagus nerve (cranial nerve X) innervates pharyngeal arch-derived muscles of the head and neck as well as providing presynaptic parasympathetic innervation of visceral targets throughout the thoracic and abdominal cavities^83^. In mammals, vagal motor (mX) neurons are organized into two anatomically distinct nuclei: the nucleus ambiguus, which mostly contains branchiomotor neurons innervating pharyngeal arch-derived muscles, and the dorsal motor nucleus, which contains parasympathetic visceral motor neurons^84,85^. In zebrafish, and in mammalian embryos^86^, both populations reside in a single elongated (∼170 µm long) motor nucleus in posterior hindbrain rhombomere 8 (r8). mX neurons are among the last cranial motor neurons to arise, first appearing in ventral r8 in zebrafish at 24 hpf and migrating dorsally to include about 350 motor neurons by 3 dpf. Within this nucleus, anterior-posterior mX neuron topography is conserved: anterior mXns innervate pharyngeal arch (PA)4-PA7 muscles and posterior mXns innervate the viscera (Fig. 5 A,B)^36,87^. Furthermore, motor neurons anterior and medial to the mX nucleus contribute to the glossopharyngeal nerve (CN IX) which innervates PA3 muscles (Fig. 5C; Movie M2). In mammals, mIX neurons similarly lie at the anterior end of the nucleus ambiguus.

**Fig. 5:**
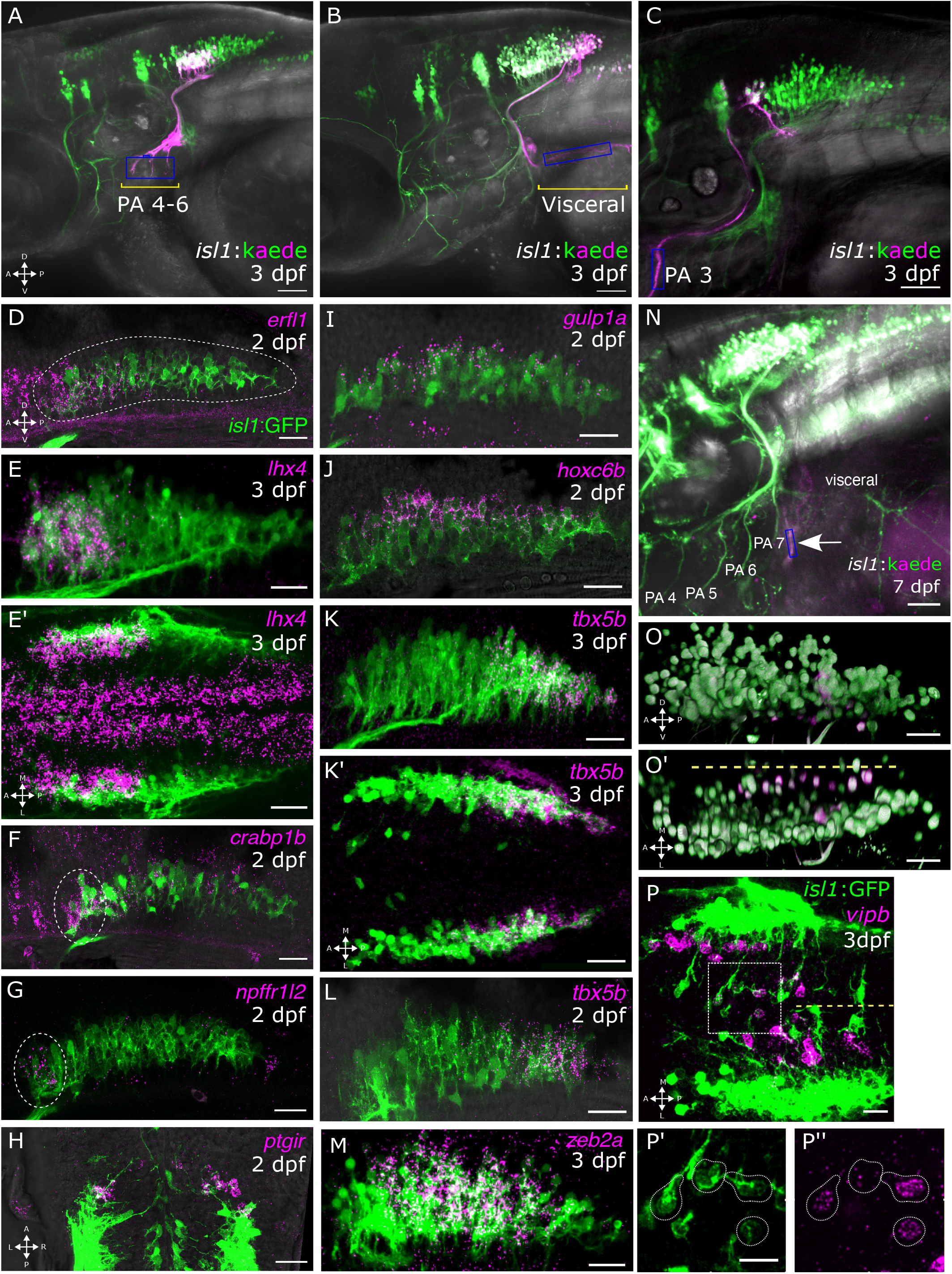
Vagus motor neuron topography corresponds to distinct transcriptional domains. **(A-C)** Retrograde labeling of mX and mIX neurons at 3dpf. Lateral views, with photoconverted Kaede in magenta, unconverted Kaede in green. Blue boxes, photoconversion ROI. (A) Photoconversion of the PA 4-6 branches labels neurons in the anterior half of the mX nucleus. (B) Photoconversion of the visceral branch labels neurons in the posterior half of the nucleus. (C) Photoconversion of PA3 branch labels mIX neurons at the anterior end of the nucleus (see supplemental movie M1). **(D-M)** HCR *in situ* validation of mX genes; HCR signal in magenta and Tg(*isl1*:GFP) in green. Images are of lateral views of the mX nucleus with anterior to the left and dorsal to the top, except E’, K’ and H where the axes are shown in the panels). **(N-O’)** Retrograde labeling of the putative heart-innervating vagal neurons. (N) Lateral view of a 7dpf Tg(*isl1*:Kaede) embryo photoconverted from the cardiac sub-branch (arrow; blue box indicates the photoconversion ROI); (O, O’) IMARIS rendering of the neurons backfilled in N (magenta). Lateral (O) and dorsal (O’) views, showing a medial longitudinal column distinct from the muscle- and viscera-innervating populations. Yellow dotted line in O’ marks the midline. **(P)** *vipb* HCR at 3dpf, dorsal view with anterior to the left. Yellow dotted line marks the midline. (P’,P’’) Magnification of inset in P, showing the Tg(*isl1*:GFP) (P’) and *vipb* HCR channels separately (P’’). HCR panels are maximum-intensity projections of ∼20µm z-stacks. Scale bars: 100µm (A-C, N); 20µm (D-M, O, O’, P); 10µm (P’, P’’)

In our atlas, mX neurons segregate into three distinct clusters (clusters 2, 3, and 5, Fig. S2D) corresponding to anterior, dorsal, and posterior mX neurons, respectively, based on HCR validation (summarized in Fig. S5A). Cluster 2 (Fig. S2D) corresponds to neurons at the anterior end of the mX nucleus. Top DEGs of this population include *erfl1*, encoding a transcription factor; *crabp1b,* a retinoic acid binding protein, and *vwc2l*, a synaptic protein (Fig. 5D,F and data not shown). Although not a DEG due to its expression in other cranial motor nuclei, the motor neuron determinant *lhx4* is also strongly enriched in these neurons and excluded from posterior mX neurons (Fig. 5E-E’). The mX *lhx4* expression domain corresponds closely with the pharyngeal muscle-innervating neurons identified by retrograde labeling (Fig. 5A). Unlike spinal motor neurons, which require *lhx3* and *lhx4* for their normal development^88^, branchiomotor neurons express only *lhx4*, consistent with their distinct developmental requirements^88,89^. However posterior mX neurons express neither *lhx3* nor *lhx4* (Fig. 5E’, S5E-F)

We subclustered cluster 2 into mIX and mXa populations (Fig. 1D; Table 5). We found that *ptgir* and *npffr1l2 are* differentially expressed in mIX when compared to mXa. Both genes are expressed in motor neurons whose position corresponds to mIX neurons (Fig. 5G,H, single z-slices, Fig. 5C, Movie M1). Thus, although spatially continuous with mX neurons, we can distinguish mIX neurons by their distinct gene expression.

Our atlas also revealed a cluster of cells corresponding to dorsal mX neurons (cluster 3, Fig. S2D, equivalent to cluster mXd in Fig. 1B). We validated several top markers of this mXd cluster, including *gulp1a,* which encodes a phagocytosis adaptor protein, and *plcb3,* a phospholipase. Both are transiently expressed dorsally in the mX nucleus at 48 hpf (Fig. 5I and not shown). The top DEG that we validated for the mXd cluster is a hox gene, *hoxc6b* (Fig. 5J). This dorsally restricted expression of *hoxc6b* is unexpected and represents a possible D-V patterning role for a Hox factor, which canonically specifies A-P differences^11^. It remains to be determined whether the dorso-ventral differences amongst mX neurons that we detect at the transcriptional level correspond to functional differences in terms of muscle and/or organ targeting.

DEGs of posterior mX neurons (cluster 5 in Fig. S2D) include *hoxb5a*, which we previously identified as a determinant of posterior r8 that is sufficient to drive mX axons to posterior targets^36^. We validated several other DEGs, including *tbx5b*, a T-box transcription factor whose expression domain correlates closely with viscera-innervating mX neurons (Fig. 5K,K’, Fig. 5B). Tbx5b is expressed in mXp neurons at 2 dpf, when mXp neurons are already committed to visceral innervation^36^ but before their axons have extended to the viscera (Fig. 5L), suggesting that the parasympathetic function of mXp neurons may be under early genetic control.

A previously uncharacterized vagal sub-branch extends posterior to the muscle-innervating PA7 branch and moves in time with the heart (Fig. 5N, movie M2). Photoconversion of Tg(*isl1*:Kaede) in this branch retrogradely labels a population of mX neurons that are distinct from either the muscle- or viscera-innervating populations described above (Fig. 5O, O’). These neurons lie medial to the main vagus motor nucleus in a distinct longitudinal column (Fig. 5O’). Subclustering of cluster 5 identified a subtype, mXm (for medial mX) (Fig. 1B, Table 6), consisting mostly of cells from embryos older than 48 hpf, which differentially expresses *nr4a2b*, an orphan nuclear receptor, and *vipb*, a neuropeptide. By HCR, these genes are selectively expressed starting at 3 dpf in medial r8 Tg(*isl1*:GFP)^+^ neurons and not elsewhere in the vagus nucleus (Fig. 5P-P’’, S5G-G’’’). Based on their position, axon projections, and *vipb* expression, we hypothesize that these neurons are preganglionic projections to the intracardiac ganglia (see Discussion).

We also identified pan-vagus motor neuron genes *zeb2a*, a zinc-finger E-box binding transcription factor^90^, (Fig.5M) and *tmeff2b,* a transmembrane protein^91^. Tmeff2b is initially expressed across the vagus nucleus at 48 hpf, becomes posteriorly biased by 3dpf, and is restricted to a discrete medial A-P domain by 6dpf (Fig.S5 B-D’). Both genes are statistically enriched in mXd, which spans the whole A-P axis of the vagus motor nucleus, and is the only population to represent anterior, medial and posterior mX neurons.

These results show that the zebrafish vagus motor nucleus, though a single continuous anatomical structure, is transcriptionally heterogeneous, comprising five molecularly distinct classes (Fig. S5A): anterior-most mIX, anterior mX, dorsal mX, posterior mX and medial mX, each of which has a distinct peripheral target as defined by retrograde labeling.

### Similarity of cranial motor neuron transcriptomes in fish and mammals

Cranial motor neuron organization is a deeply conserved feature of vertebrate brains, although the positions and functions of the muscles and organs that individual cMNs innervate have diverged^1,3^. To investigate whether the molecular differences between and within the cranial motor nuclei that we identified in zebrafish are similar to those in mammals, we performed a cross-species analysis between our atlas and cMNs extracted from a large-scale whole embryo mouse development sci-RNA-seq atlas^19^. Mouse motor neurons sampled between E8.0 and E15 cluster into cMNs (*Tbx20*+) and spMNs (*Mnx1*+) like in zebrafish (Fig. S6A, B; Suppl. Fig. 6C). cMNs were further subsetted and re-processed, generating an atlas of developing mouse cMNs composed of 16,309 cells and 8 clusters (Fig. 6A, B).

**Fig. 6:**
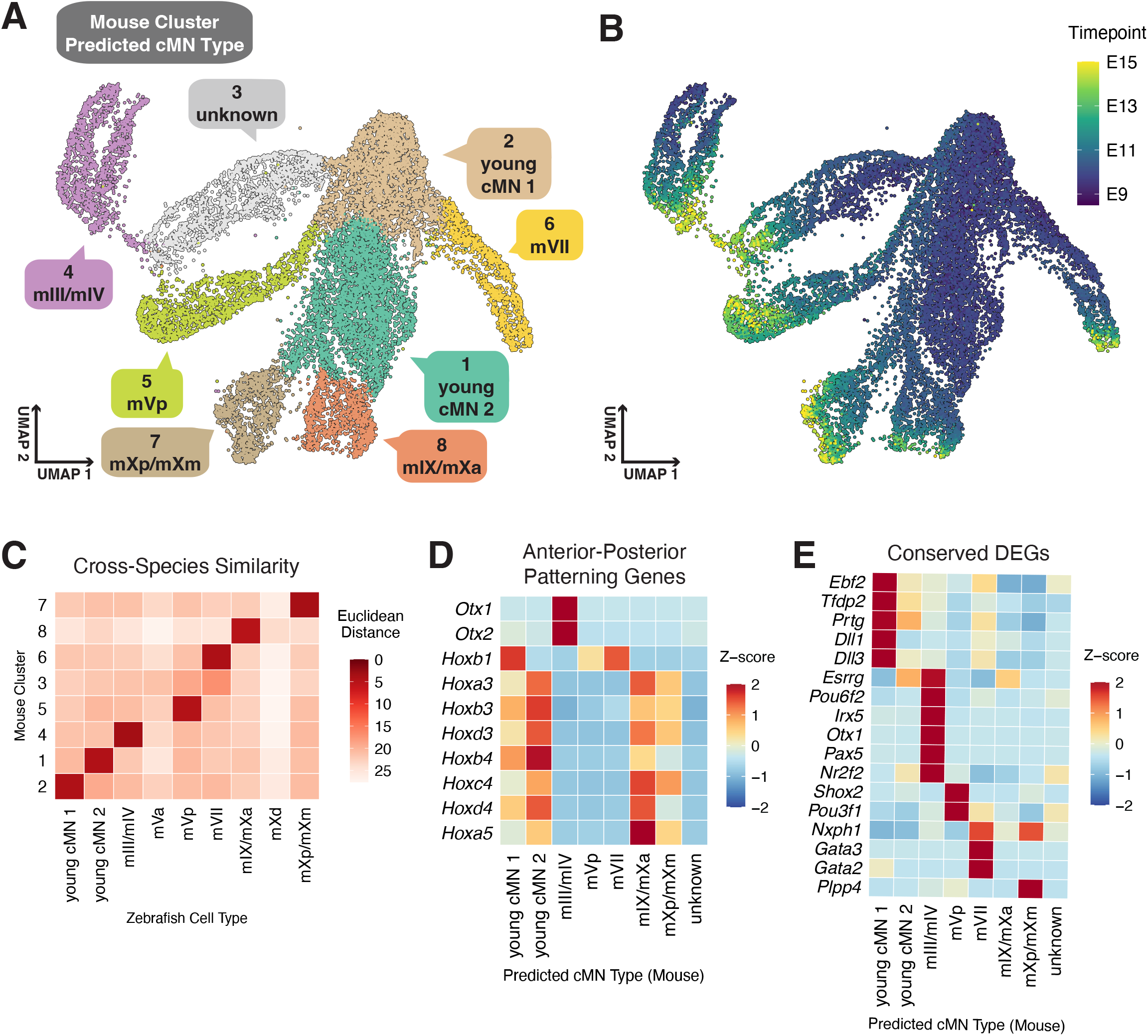
Cross-species analysis of zebrafish and mouse cranial motor neurons. **(A)** UMAP of mouse cMNs. Colors correspond to zebrafish clusters shown in Fig. S2D). Labels indicate the mouse cluster number (top) and the SATURN-predicted cMN type (bottom). **(B)** UMAP of mouse cMNs colored by the timepoint in embryonic days (E) at which cells were sampled. **(C) H**eatmap comparing our initial 9 zebrafish clusters (named by the cMN types they represent) with the eight mouse cMN clusters. Darker red colors indicate shorter Euclidian distances. **(D)** Heatmap of Otx- and Hox-family gene expression across the mouse cMN clusters, supporting the predicted identities (e.g., Otx1/Otx2 in mIII/mIV, Hoxb1 in mVII, and posterior Hox genes in mIX/mX clusters. Otx- and Hox-genes were first filtered for a minimum expression in at least 10% of any mouse cluster. **(E)** Heatmap of zebrafish cluster DEGs from Table 1. Table 1 genes were converted to mouse orthologs, filtered for a minimum expression in at least 10% of any mouse cluster, then filtered for genes that are differentially expressed in the corresponding cMN type.

We integrated our zebrafish and mouse cMN atlases using SATURN, a deep learning-based method that uses protein language models to group functionally related genes from different species into embeddings, which are then coupled with gene expression^92,93^. Cranial motor nuclei annotated from the original 9 zebrafish clusters (Fig. S2D) were used as cell type labels to ensure sufficient coverage of each nucleus. We predicted the identity of the mouse cMN clusters by computing 50 principal components (PCs) of the SATURN joint embedding, then calculating the Euclidean distance between the centroids of the mouse clusters and zebrafish cMN types in PC space. The majority of the mouse cMN clusters exhibited a one-to-one correspondence with zebrafish cMN types (Fig. 6C), enabling us to predict the identity of the mouse clusters accordingly (Fig. 6A).

The predicted cMN types in mouse were supported by the expression of anterior-posterior patterning genes. *Otx1* and *Otx2,* both midbrain determinants, are enriched in the mouse cluster assigned by SATURN as mIII/mIV; *Hoxb1*, a determinant of hindbrain rhombomere (r)4 identity, is enriched in the mouse cluster assigned as mVII; while Hox-3, Hox-4 and Hox-5 genes, which are expressed in the posterior hindbrain, are detected in mouse clusters assigned as mIX and mX which lie in r7 and r8 (Fig 6D).

We mapped the mouse orthologs of the top 20 DEGs from each of our zebrafish clusters (Table 1) onto the mouse clusters. While many of these genes are either not detectable or not expressed in the corresponding mouse clusters, some are (Fig. 6E). Interestingly, the process of selecting orthologous genes that are expressed in corresponding fish and mouse clusters enriched for transcription factors and developmental signaling pathway components. Based on their conserved expression, these genes are likely to be the upstream determinants of conserved aspects of specific cMN identities (Fig. 6E).

## DISCUSSION

Classical studies define cranial motor neurons (cMNs) by their rhombomere of origin and their peripheral targets^1,60^, with subsequent studies describing the general specification of cMNs as a class^1,9^. It has remained unknown whether gene expression can resolve individual nuclei and their functional neural subtypes. We address this with a developmentally resolved single-cell atlas, validated extensively by HCR. While many genes are shared across cranial motor nuclei, each nucleus (and in several cases functional subdivisions within a nucleus) has a distinct molecular signature. This study adds key developmental information to a body of work that has focused primarily on spinal MNs in adult tissue^28,30,31^.

### Multiple parameters shape cranial motor neuron identity

The molecular identity of individual cMNs is shaped by three intersecting parameters. The first is shared function: for example, the three extraocular nuclei (mIII, mIV, mVI) arise from different rhombomeres yet act together to move the eye and are more transcriptionally similar to each other than to other cMNs, suggesting that functional similarity and common synaptic targets are promoted by a common program. The second parameter is rhombomere of origin: for example, in the trigeminal nucleus, r2-derived (mVa) and r3-derived (mVp) neurons differ markedly in gene expression and muscle targets, dependent on upstream determinants of rhombomere identity. The third parameter is developmental time: for example, in the facial nucleus (mVII), gene expression changes as neurons migrate caudally from rhombomere 4, where transcriptional states change with maturation and cell position.

### Molecular identity precedes and predicts connectivity

Transcriptional identity precedes and predicts peripheral connectivity. In the vagus nucleus (mX), posterior, visceral projecting neurons selectively express *tbx5b* at 2dpf before their axons extend toward visceral targets, implying early genetic commitment to a visceral fate. Molecular subtypes also map onto functional groups defined by retrograde labeling. Trigeminal mVa and mVp neurons innervate different first-arch muscles, and a dorsolateral *sim1a/opn8b*-positive oculomotor subset corresponds to early-born neurons innervating the medial rectus implicated in convergent eye movements^48,49^. These genetic programs are thus in place before circuit assembly.

### Cranial MNs heterogeneously implement a modified spinal MN developmental program

Cranial motor neuron identities are specified through a modified spinal motor neuron program. Lhx3 and Lhx4 are established determinants of ventrally-projecting motor neuron identity in the spinal cord^88,89,94,95^. In the chick brainstem, Lhx3 is restricted to the accessory abducens and hypoglossal nuclei and is absent from branchiomotor and visceromotor populations^88,96^. However, Lhx4 expression in vagal motor neurons has not been directly examined in any species^12^. We show that the zebrafish vagus nucleus is heterogeneous along the A-P axis: anterior, branchiomotor-like mX neurons express *lhx4* but not *lhx3*, while posterior, viscera-projecting neurons express neither (Fig. 5, S5E-F). Like Lhx4, many markers reported for individual cranial nuclei are in fact shared across several cMN populations^66,85,97^. This atlas allows us to separate genes that confer nucleus or subnucleus identity from broader motor neuron identity programs.

### Molecular identification of novel cranial motor neuron populations

Our atlas resolves both functionally divergent, and closely related cMN populations. Glossopharyngeal (mIX) neurons, though continuous with the anterior vagus (Movie M1), are molecularly distinct. This may reflect retinoic acid (RA) responsiveness, as mIX neurons are relatively RA-insensitive^98^, and express *crabp1b* and other RA modulating factors (Fig. 5F, Table 5). We also identify a novel mXm cluster defined by vipb, representing a fourth class of mX neuron. Several observations suggest that these neurons might serve a cardiac role: 1) they are retrogradely labeled from a putative heart-innervating mX branch; 2) they express vipb, whose ortholog, VIP, is released by vagal efferents to regulate heart rate in mammals^99–102^. 3) In zebrafish, vagal stimulation evokes bradycardia, with the heart becoming responsive to cholinergic stimulation at 5dpf and intracardiac ganglia adjacent to the sinoatrial region receiving vagal input^103,104^. The *vipb*-expressing mXm population is therefore a strong candidate for preganglionic parasympathetic input to the zebrafish intracardiac ganglia, analogous to cardiac-projecting mammalian DMV neurons.

### cMNs use a conserved developmental program

Several aspects of the cMN organization we describe are conserved in mouse. Cross-species comparison with cMNs from a whole-embryo mouse dataset predicts a largely one-to-one correspondence between zebrafish and mouse nuclei, despite the divergent muscles and organs that they control, implying that the transcriptional logic distinguishing cMN identities is deeply conserved. For most clusters, the similarities that underlie this correspondence are deep and dispersed in the transcriptomic data, rather than being driven by a small number of orthologous genes, and in two cases, cell types that are transcriptionally very distinct in the zebrafish dataset (mVa and mXd) are not resolvable in the mouse dataset even though there are homologous neurons in mammals. On the other hand, the presence of a cluster in the mouse dataset that has no zebrafish correlate (cluster 3, Fig. 6A) may reflect the fact that mammals have two cranial motor nerves that zebrafish lack (CN XI and XII). The transcription factors we have identified as being similarly expressed in zebrafish and mouse cranial motor neuron populations are top candidates for being determinants of conserved aspects of motor neuron identity.

### Outstanding questions and disease relevance

Our atlas defines a set of testable questions. Most immediate is whether the nucleus and subtype-specific genes we identify are required for cMN fate specification and axon targeting. For example, mutational analysis can determine whether transcription factors expressed in muscle-versus viscera-targeting mX neurons are required to specify those functional differences. Retrograde labeling can address whether new populations identified by gene expression here, such as mXm and mXd, correspond to a distinct target group, while tracing and perturbation of the *vipb-*expressing mXm neurons should establish whether they regulate heart rate. Knock-in reporters will enable live imaging of neurons generated at different stages of mVII neuron migration, to address how developmental timing determines OEN versus FBMN identity. Finally, our discovery of distinct, early mVa versus mVp markers provides tools to revisit the role of Hox genes in specifying cranial motor neuron identity.

Defects in cMN specification, migration and axon guidance underlie human congenital cranial dysinnervation disorders and contribute to congenital central hypoventilation syndrome^5,6^. Our results provide a basis for understanding how genetic perturbations selectively disrupt individual cranial motor pathways or instruct upstream circuit connectivity with higher-order brain centers, particularly given the apparent deep transcriptional similarity of zebrafish and mouse cMN subtypes. Our results show that transcriptional identity alone is sufficient to define individual nuclei and their functional subtypes within a closely related, yet functionally diverse population of motor neurons.

## MATERIALS AND METHODS

### Embryo dissociation, flow cytometry and sequencing

Tg*(isl1*:Kaede) embryos were raised to 24, 48, 72 or 144hpf respectively prior to being anesthetized in 400mg/L tricaine methanesulfonate (MS-222). Hindbrains were dissected in Calcium-Free Ringers / EDTA / 10% MS-222, prior to dissociation to single-cell suspension by pipetting in 0.25% trypsin-EDTA for 5 minutes. Cells were briefly spun down and transferred to dPBS + 3% BSA. Prior to FACS, cell health was evaluated using 2ug/mL DAPI. Kaede^+^ cells were isolated on a BD FACS ARIA II flow cytometer using a 100uM nozzle and collected into dPBS + 2% BSA. Collected cells were concentrated by centrifugation at 300 x g for 5 minutes at 4°C and resuspended to an optimal concentration of 1,200 cells/uL. Cell viability was assessed using a TC20 Automated Cell Counter. Libraries were prepared using the 10X Genomics Chromium 3’ mRNA v3 kit, prior to sequencing on a Novaseq S1 Flowcell. 10X libraries were sequenced to an average depth of 44,057 reads per cell (2,015 genes per cell).

### Sequencing, read processing, and cell filtering

Following sequencing, the coding sequence for Kaede (https://zfin.org/ZDB-EFG-080131-2#summary) was appended to the “Danio_rerio.GRCz11.105.gtf” (https://ftp.ensembl.org/pub/release-105/gtf/danio_rerio/), (filtered for protein coding sequences using CellRanger “mkgtf”) and “Danio_rerio.GRCz11.dna.primary_assembly.fa” files from Ensemble (https://ftp.ensembl.org/pub/release-105/fasta/danio_rerio/dna/), and a custom reference was assembled using the CellRanger-7.0.0 “mkref” function (default settings). Reads were aligned using CellRanger “count” (default settings).

For each 10x run, low quality cells with less than 500 UMIs, greater than 3 standard deviations above the mean UMI, and/or percent mitochondrial reads greater than 25% were removed.

### Single cell RNA-seq analysis

Data were processed using the Monocle3 (develop branch, v1.4.16) using defaults except where specified: *align_cds(residual_model_formula_str = “∼Size_Factor”)* and *reduce_dimensions(cds, preprocess_method = “Aligned”).* All analysis code, including UMAP and clustering parameters for each step are available in the github repository.

Trapnell Lab (sci-RNA-seq): We first subsetted the motor neuron projection groups from each published dataset^26,43,44^. Cells from all perturbations were retained. The *tbx20+ mnx1-* cluster was selected for further analysis.

Moens Lab (10x): Cells were then processed, and a *tbx20+ isl1+ chata+* cluster was selected for further analysis.

Farrell Lab, Daniocell (10x): The Daniocell dataset was downloaded from https://www.ncbi.nlm.nih.gov/geo/query/acc.cgi?acc=GSE223922 and https://daniocell.nichd.nih.gov/ and loaded using Seurat (v.5.1.0). We selected cells labeled “glia_11” and constructed a Monocle3 cds. First, gene names were converted to GRCz11 ENSEMBL gene names using the Lawson Lab (v4.3.2) gene annotations. LOC-, XLOC-, NC-genes, and genes with no associated ENSEMBL gene name were removed. Counts from genes with the same ENSEMBL gene name were consolidated by summing counts. Cells were processed and a *tbx20+ isl1+ chata+ mnx1-* cluster was selected for further analysis.

Shendure Lab (sci-RNA-seq): The mouse dataset was downloaded from https://cellxgene.cziscience.com/collections/45d5d2c3-bc28-4814-aed6-0bb6f0e11c82 (CZ CELLxGENE dataset ID: e6be6391-47a9-4a22-925d-27596e475318). We specifically downloaded “Major cell cluster: CNS neurons” and loaded the dataset into Scanpy (v1.10.2). Cells labeled “cranial motor neuron” and “spinal cord motor neuron” in the “cell_type” metadata column were selected for generation of a Monocle3 cds. Cells were subsetted to stage E15 and earlier and processed. A *Tbx20+ Mnx1-* cluster was subsetted and processed. We removed a Chat-Slc17a6+ cluster, which we believe are not motor neurons, and reprocessed the cells. Marker genes were computed using the Monocle3 *top_markers()* function, significance determined by a two-sided likelihood test with multiple testing correction.

### Integration of zebrafish cMN single-cell datasets

Datasets were converted to Seurat objects and processed using *NormalizeData()* and *FindVariableFeatures(nfeatures=2500).* The datasets were integrated using *FindIntegrationAnchors(dims = 1:30, k.filter=200)* and *IntegrateData()*. The resulting integrated counts matrix was exported to generate a Monocle3 cds for processing. UMAP coordinates and clusters computed with the integrated counts matrix were combined with the raw counts matrix for further analysis.

### Differential gene expression analysis

Cranial versus spinal motor neuron comparison: Expression values were first aggregated for each embryo across clusters into ‘pseudo-cells’. Differential gene expression between the clusters was performed using a generalized linear regression model using the Monocle3 *fit_models()* function.

Identification of cluster and nuclei markers in the integrated zebrafish dataset: To account for the significant technical differences between the datasets (most notably, sample size and UMI counts), cluster markers were identified using a cross-dataset consistency approach. Within each dataset, counts were normalized to 10,000 counts per cell and log transformed. Log2 fold-changes were computed for each gene by comparing normalized expression in cells within versus outside each cluster. A gene was retained as a marker if, across at least two of the datasets, it was (1) detected in >=10% of cells in the cluster and (2) exceeded log2FC>1. Markers were ranked according to the median log2FC across datasets and detection ratio (fraction of cells in the cluster expressing the gene versus cells outside the cluster).

### Cross-species analysis

To account for substantial UMI count differences between the zebrafish sci and 10x datasets, raw expression values of 8 random cells within each motor nucleus of the Trapnell sci dataset were aggregated into ‘pseudo-cells’. The integrated zebrafish and mouse datasets were exported to AnnData objects using Scanpy (v1.10.2) and used to train SATURN with motor nuclei labels for zebrafish and cluster labels for mouse, *num_macrogenes=2000* and *hv_genes=8000*, and ESM2-derived protein embeddings for each species. The outputs were processed in Monocle3 with *preprocess_cds(num_dim=50, norm_method=“none”)* and cross-species cell type similarity scores were determined by calculating Euclidean distances between the centroids of zebrafish and mouse clusters.

### Data and code availability

Raw data will be available upon publication: NCBI BioProject PRJNA1470670. Project information and processed CellRanger outputs can be found under NCBI GEO GSE336035. Code used to analyze the single-cell data and generate plots in the figures is available at https://github.com/rikuyasutomi/cranial-motor-neuron-atlas.

### Retrograde labeling

Tg(*isl1*:Kaede) embryos were collected and incubated at 28.5°C. At 72hpf animals were anaesthetized with MS-222 and mounted in 1.2% low-melt agarose. Axon branches innervating described structures were photoconverted by 405 illumination (Zeiss LSM 700), and > 1 hour was allowed for diffusion of converted protein into cell bodies prior to follow-up imaging on a Stellaris 5 DMI 8 confocal microscope with a 40x water immersion objective, 1-2µm z-stacks.

### HCR sample prep

Zebrafish embryos were staged according to needed timepoint before being anesthetized by MS-222 (0.4% solution) and fixed in 4% PFA / 4% sucrose solution overnight at 4°C. Following fixation, embryos were washed into MeOH and stored at −20°C until use. Prior to HCR, embryos were rehydrated through a MeOH/PBST gradient. HCR v3.0 (Molecular Instruments) was performed as previously described^37^ with adjustments to proteinase K incubation time based on age (28 hpf: 5-minute incubation, 48 hpf: 20 minutes, 3 dpf: 30 minutes, 6 dpf: 60 minutes). HCR experiments used buffers and amplifiers from Molecular Instruments. HCR v3.0 probe pairs were either ordered through Molecular Instruments, Inc. or designed in-house^105^. Following the completion of HCR v3.0 protocol, embryos were washed through a glycerol / PBS gradient. Samples were stored in 75% glycerol / 25% PBS at 4°C until imaging. Embryos were dissected to isolate the brain and mounted either with the ventral or the lateral surface on a 24 x 60 mm coverslip in 75% glycerol. Another 18 x 18 mm coverslip was placed on top with posts of high vacuum grease to prevent the brain from being flattened. We imaged the mounted embryos on a Stellaris 5 DMI 8 confocal microscope with a 40x water immersion objective, 1µm z step size. Images were processed using Fiji/ImageJ. HCR validation of genes not shown in this paper are available on the Zebrafish Information Network gene expression database.

### Zebrafish care and maintenance

*Danio rerio* animals were raised at the Fred Hutchinson Cancer Center facility in accordance with IACUC-approved protocols. All experiments were carried out in accordance with IACUC standards. Fish were bred and maintained according to standard protocols^106^. Sex is not a relevant biological variable in our experiments, as they are carried out before sex is determined in zebrafish^107^. Transgenic lines used in this study include Tg*(isl1*:GFP-CAAX)fh474^36^ (referred to throughout as Tg(*isl1*:GFP)), Tg(*isl1*:Kaede)ch103^36^*, T*g(*Tcf21*:mCherry)pd108^108^, Tg(*Olig2*:Gal4) (gift of Karina Yaniv), and Tg(UAS:Kaede)s1999t^109^.

## Acknowledgements

We thank Jason Stonick, Veronica Sikora, Haley Dean, Jon Cooper and Trapnell lab members, for their technical and intellectual support during this work and for critical reading of the manuscript; Chengxiang Qiu for data sharing and support with the mouse data; Sanjay Kottapali and Wei Yang for technical help with the cross-species analysis; The Fred Hutch Comparative Medicine Core, particularly Amelia Bothell, Nick Arbasak and Ben Zuidema for zebrafish care; and the Fred Hutch Genomics Core, particularly Dolores Covarrubias for help with the 10X preps.

## Author contributions

ID, AQS, RY and CBM designed and performed HCR experiments and data analysis. RY and AQS performed computational analysis. AQS and AJI performed 10X sequencing experiments. AQS, RY and CBM wrote the manuscript. All authors edited the manuscript. CBM and CT provided supervision and funding acquisition.

---

**Fig. S1:**
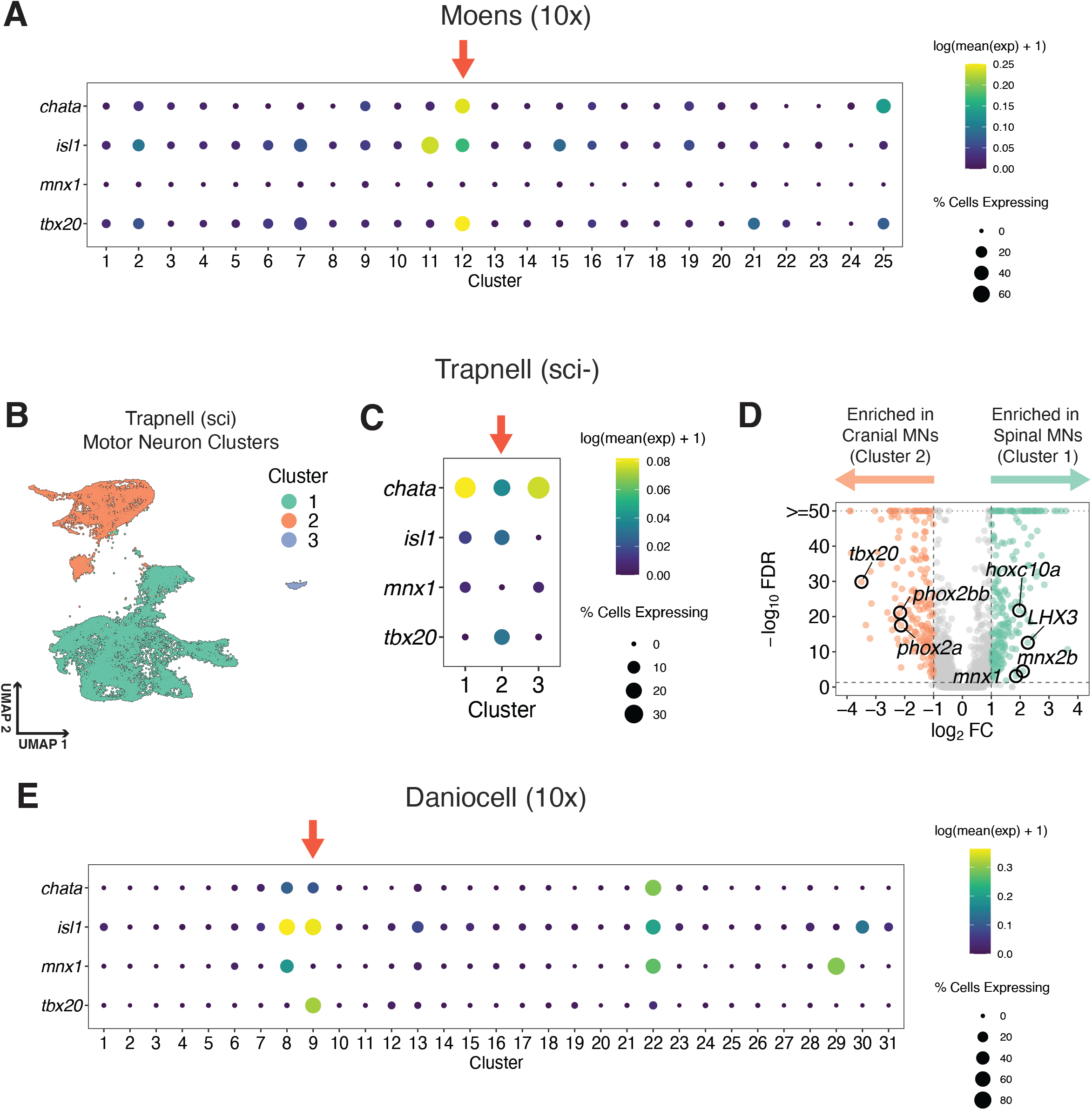
Identification of cranial motor neurons in each single-cell dataset. **(A)** Dot plot of *chata, isl1, mnx1, and tbx20* expression across all clusters computed in the Moens 10x dataset. The red arrow indicates the cluster we identified as cMNs.**(B)** Two-dimensional UMAP embedding of the Trapnell sci-RNA-seq motor neurons, colored by cluster. **(C)** Dotplot of *chata, isl1, mnx1, and tbx20* expression across the three Trapnell sci-RNA-seq clusters. The red arrow indicates the cluster we identified as cMNs. **(D)** Volcano plot of DEGs between Trapnell sci-RNA-seq cluster 2 (cMNs) and cluster 1 (spMNs). A few genes known to distinguish cMNs from spMNs are labeled. **(E)** Dotplot of *chata, isl1, mnx1, and tbx20* expression across all Daniocell 10x clusters we computed from the “glia” group. The red arrow indicates the cluster we identified as cMNs.

**Fig. S2:**
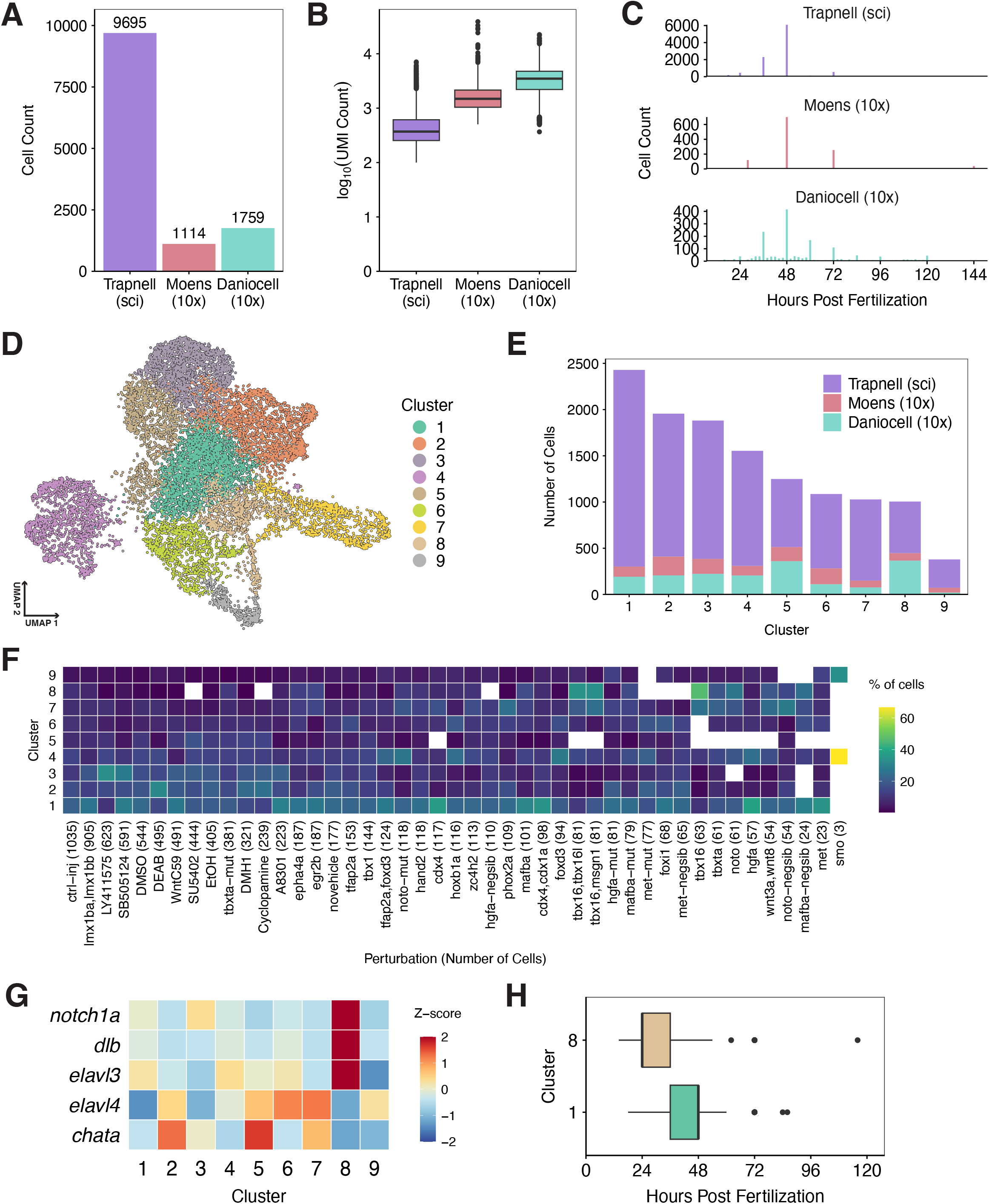
Generation of an integrated cranial motor neuron single-cell atlas. **(A-C)** Comparison of cell counts (A), Unique molecular identifier (UMI) counts (B), and distribution of sampling timepoints (C) between the three datasets. (D) Two-dimensional UMAP embedding of the integrated atlas colored by clusters computed from the corrected (integrated) expression matrix. (E) Number of cells in each cluster and the composition by dataset. (F) Heatmap representing the distribution of cells across the 9 clusters for each perturbation in the Trapnell sci-RNA-seq dataset. The color scale represents the percent of cells, for a given perturbation, that contribute to each cluster. Empty boxes indicate zero cells, which increase in number for perturbations that contributed very few cells overall to the dataset. No individual perturbation appreciably altered the distribution of cells across cMN clusters. (G) Heatmap of genes distinguishing immature (*notch1a, dlb,* and *elavl3*) from more mature (*elavl4* and *chata*) cMN states. (H) Comparison of sampling timepoints between clusters 1 and 8.

**Fig. S3:**
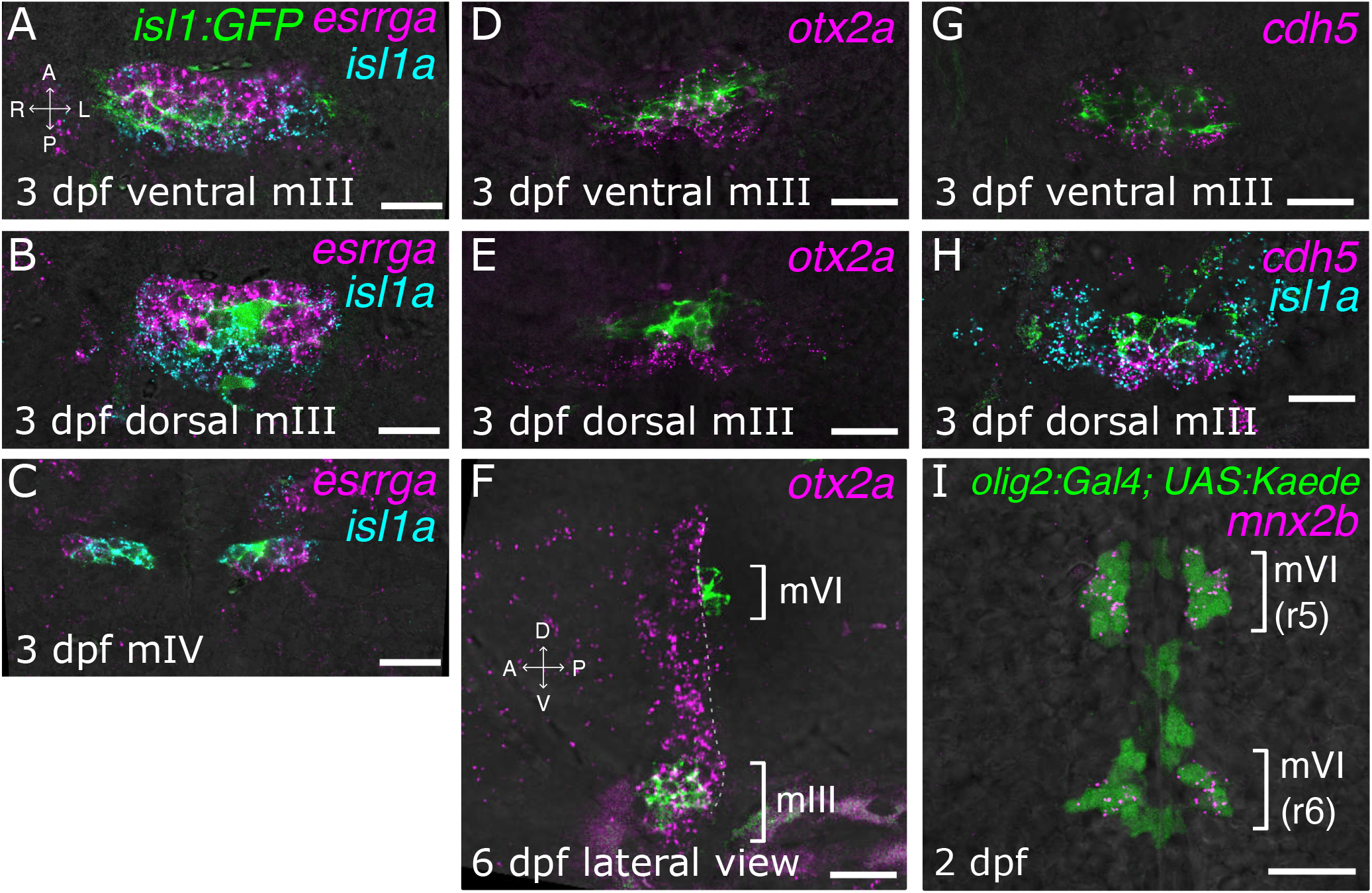
Further validation of extraocular motor neuron markers. HCR in situ validation of gene expression in eye-innervating motor neurons. mIII: oculomotor; mIV: trochlear; mVI: abducens motor nuclei. HCR products are magenta or cyan, as noted; green is Tg(*isl1*:GFP) except in I where green is Tg(*olig2*:Gal4; UAS:Kaede). All images are single z-sections in ventral view with anterior to the top except F which is a lateral view as indicated by axes in the panel. Where noted, “ventral” and “dorsal” z-sections are ∼20 apart, to capture the dorso-ventral depth of mIII and mIV nuclei. **(A-C)** *esrrga* (magenta) and isl1a (cyan) expression in mIII and mIV neurons at 3dpf. **(D-F)** *otx2a* expression in mIII neurons at 3 dpf. (D,E) ventral (D) and dorsal (E) mIII planes at 3dpf. (F) *otx2a* at 6dpf, lateral view, showing persistent mIII expression. Dotted line marks the posterior limit of *otx2a* expression domain corresponding to the midbrain-hindbrain boundary.**(G-H)** *cdh5* (magenta) and isl1a (cyan in G) expression in mIII at 3dpf, ventral (G) and dorsal (H) planes. (I) *mnx2b* expression (magenta) in hindbrain mVI neurons marked by Tg(*olig2*:Gal4; UAS:Kaede) (green). Scale bars = 20µm.

**Fig. S4:**
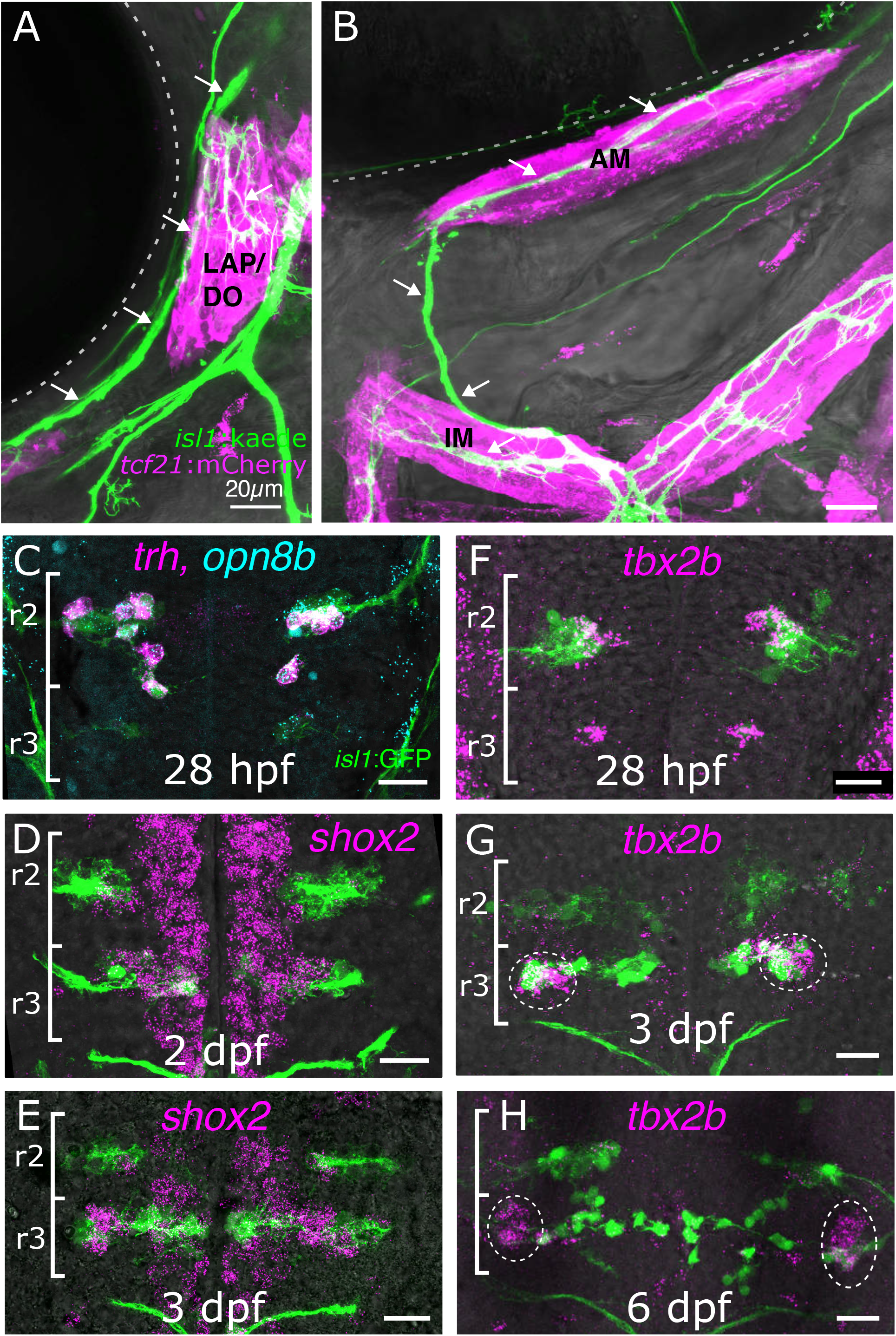
Trigeminal motor neuron (mV) targets and further validation of mVa- and mVp markers. **(A-B)** lateral (A) and ventro-lateral (B) views of mV axons Tg(*isl1*:Kaede); green) innervating first pharyngeal arch muscles (TgBAC(*tcf21*:mCherry-NTR); magenta) in a 4 dpf embryo. Arrows mark trigemnial motor axon tracts: the dorsal branch innervates LAP and DO muscles (A), and the mandibular branch innervates the AM muscle, and more ventrally, IM muscle (B). Dotted lines mark the eye. **(C-H)** HCR in situs validation of mV marker genes. mVa neurons are in r2; mVp neurons are in r3. All images are max-intensity projections of a ∼20-30 µm z-stack in ventral views with anterior to the top, to capture the dorso-ventral depth of mV nuclei. Green: Tg(*isl1*:GFP)); magenta and cyan: HCR signals. (C): trh and opn8b expression in mVa neurons at 28 hpf; (D,E): *shox2* expression in mVp at 2dpf (D) and 3dpf (E). *Shox2* is also detected medially in non-motor neurons along the length of the hindbrain. (F-H): *tbx2b* expression in mVp at and a subset of medial mVa neurons at 28hpf (F), and restricted to lateral mVp neurons (dashed circles) at 3dpf (G) and 6dpf (H) Scale bars = 20µm.

**Fig. S5:**
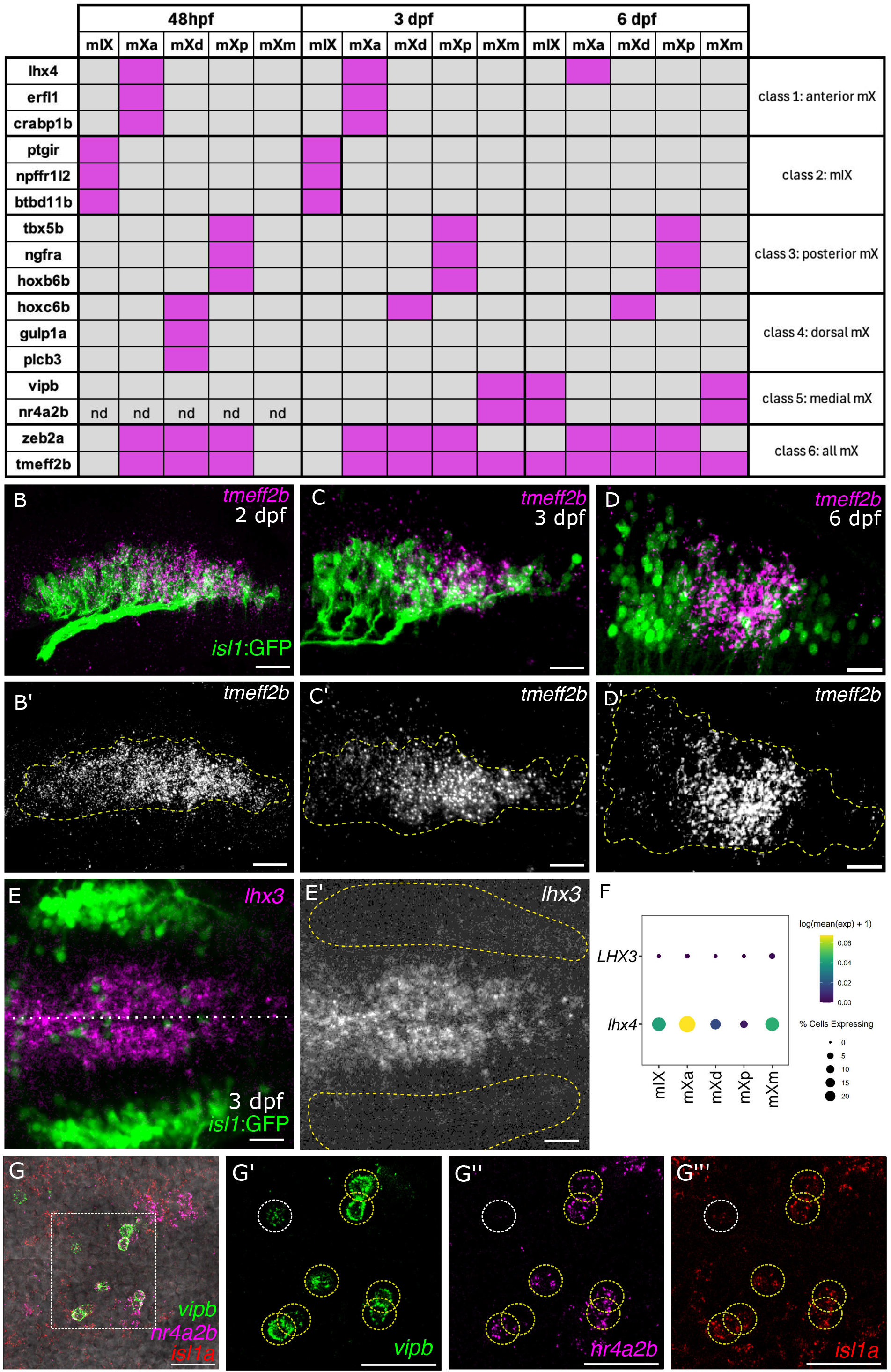
Marker genes distinguishing glossopharyngeal (mIX) and vagus (mX) motor neuron subtypes. **(A)** Summary of HCR evidence for gene expression across mIX and the mX subtypes (mXa, mXd, mXp, mXm) at 48hpf, 3dpf, and 6dpf. Magneta fill indicates expression; grey fill indicates lack of expression; nd: not done.**(B-D’)** *tmeff2b* expression (magenta (B-D) or white (B’-D’) with Tg(*isl1*:GFP) (green). Lateral views of the mX nucleus. Images are maximum-intensity projections of a ∼20 µm z-stack. (B’,C’,D’) *Tmeff2b i*s initially expressed across the nucleus with a slight posterior bias. (B,B’), becomes more posteriorly restricted by 3dpf (C,C’), and is confined to a medial domain by 6dpf (D,D’). **(E-E’)** *lhx3* (magenta (E) or white (E’) is expressed in ventral hindbrain progenitors but not in mX neurons. White dotted line in E marks the midline; yellow dots in E’ outline the mX nucleus.. **(F)** Dotplot of *lhx3* and *lhx4* expression from the single-cell data across the mIX, mXa, mXm, mXd and mXp clusters: *lhx4* is detected in mIX/mXa and mXm but not in mXd or mXp, whereas *lhx3* is absent throughout. **(G-G’’’)** (G) Triple 3dpf HCR with *vipb* (green), *nr4a2b* (magenta) and *isl1a* (red). Ventral view of a single z-section, anterior to the left. (G’-G’’’) Zoom-in of boxed region in (G). Demonstrates colocalization of all three genes in many mXm neurons (yellow dashed circles), while a subset of cells are *vipb*+ / *nr4a2b-* / *isl1a*+ (white dashed circles). Scale bars = 20µm (B-G’’’).

**Fig. S6:**
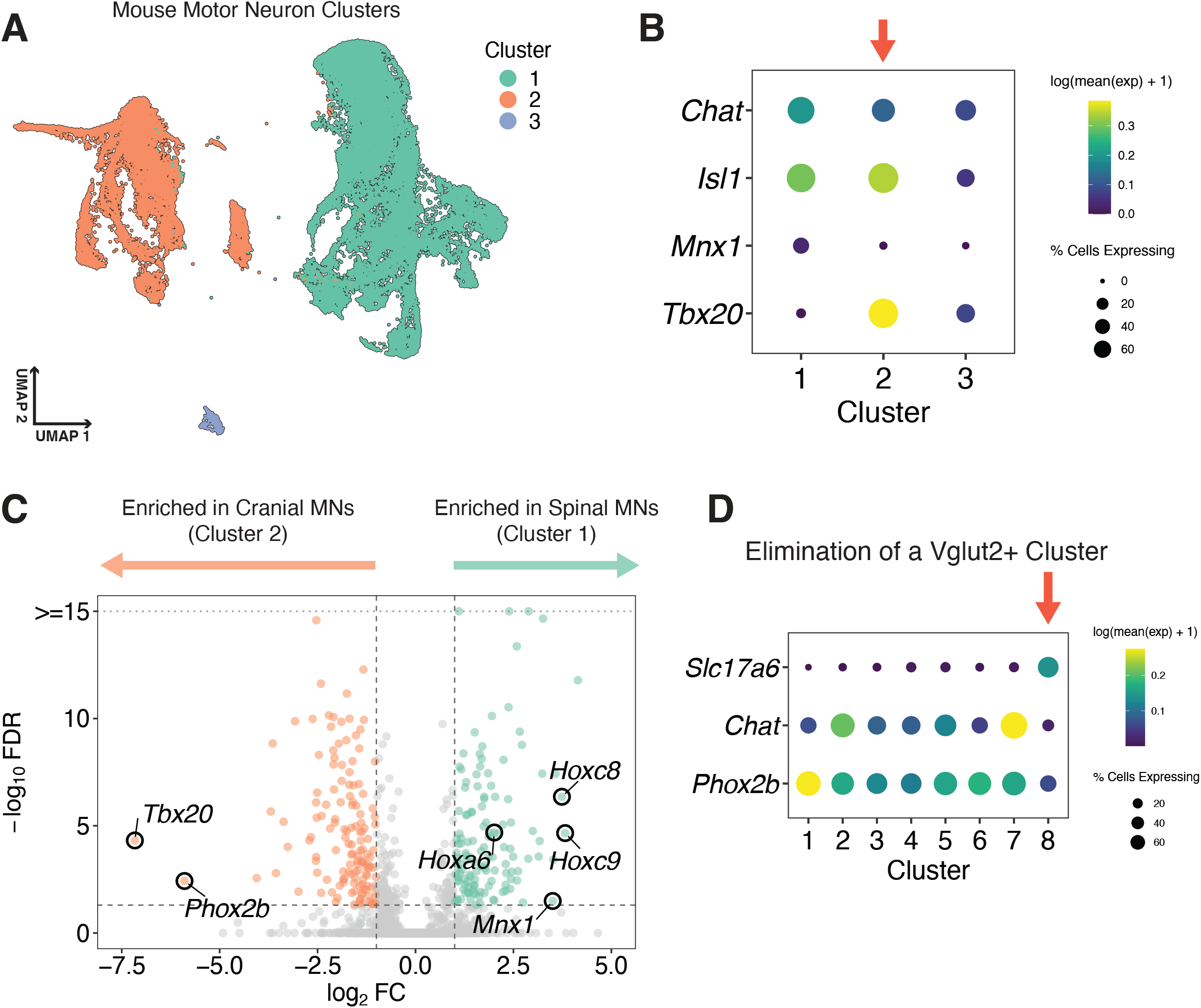
Generation of a mouse cranial motor neuron single-cell atlas. (A) Two-dimensional UMAP embedding of the mouse motor neurons, colored by cluster. (B) Dotplot of *Chat, Isl1, Mnx1, and Tbx20* expression across the three mouse clusters. The red arrow indicates the cluster we identified as cMNs. (C) Volcano plot of DEGs between mouse cMNs (cluster 2) and spMNs (cluster 1). A few genes known to distinguish cMNs from spMNs are labeled. (D) Identification of a Chat-Slc17a6+ cluster (red arrow) that we believe does not correspond to cMNs. We eliminated this cluster and reprocessed the cMNs to generate our final mouse atlas (Fig. 6A).

**Movie M1:** mIX neurons are continuous with and medial to the mX nucleus 90° Rotation from lateral to dorsal view of a z-stack showing mIX neurons labeled retrogradely from their PA3-innervating peripheral nerve (magenta, white arrow). mIX neurons form a loose nucleus with anterior-most neurons lying medially in r7 while posterior neurons are continuous with anterior-most mX neurons in r8.

**Movie M2:** Putative heart-innervating mX sub-branch moves in time with heartbeat. Movie of 4 dpf lateral view Tg(*isl1*:Kaede) (green) and *tcf21*:mCherry (magenta) across 2.2 seconds (8.6 frames/second) demonstrating the putative heart-innervating vagal motor sub-branch moving in time with heartbeat (white arrow). More anterior PA7 branch does not move in time with heartbeat (white dashed circle).

**Supplemental Table 1:** DEGs between the 9 zebrafish clusters (shown in Fig. S2D)

**Supplemental Table 2:** DEGs between the 13 annotated cMN states (shown in Fig. 1B identified in this paper based on HCR validation).

**Supplemental Table 3:** DEGs between 8 subclusters of the extraocular muscle-innervating cluster 4 in Fig. S2D identifies a population of dorsal-lateral mIII neurons (cluster 7 in this table, corresponding to mIIId in Fig. 1B)

**Supplemental Table 4:** DEGs between 3 subclusters of cluster 8 in Fig. S2D identifies a population of young mVa neurons (cluster 3 in this table, corresponding to mVay in Fig. 1B).

**Supplemental Table 5:** DEGs between 2 subclusters of cluster 2 in Fig. S2D identifies mIX neurons (cluster 2 in this table, corresponding to mIX in Fig. 1B).

**Supplemental Table 6:** DEGs between 2 subclusters of cluster 5 in Fig. S2D identifies putative heart-innervating mX neurons (cluster 2 in this table, corresponding to mXm in Fig. 1B).

**Supplemental Table 7:** DEGs between the 8 mouse cMN clusters (corresponding to Fig. 6A)

## References

1. Guthrie, S. Patterning and axon guidance of cranial motor neurons. Nat Rev Neurosci 8, 859– 871 (2007).

2. Chandrasekhar, A. Turning heads: Development of vertebrate branchiomotor neurons. Developmental Dynamics 229, 143–161 (2004).

3. Fritzsch, B. Evolution and development of extraocular motor neurons, nerves and muscles in vertebrates. Annals of Anatomy - Anatomischer Anzeiger 253, 152225 (2024).

4. Whitman, M. C. Axonal Growth Abnormalities Underlying Ocular Cranial Nerve Disorders. Annu. Rev. Vis. Sci. 7, 827–850 (2021).

5. Whitman, M. C. & Engle, E. C. Ocular congenital cranial dysinnervation disorders (CCDDs): insights into axon growth and guidance. Human Molecular Genetics 26, R37–R44 (2017).

6. Pattyn, A. et al. Coordinated temporal and spatial control of motor neuron and serotonergic neuron generation from a common pool of CNS progenitors. Genes Dev. 17, 729–737 (2003).

7. An, D. et al. Stem cell-derived cranial and spinal motor neurons reveal proteostatic differences between ALS resistant and sensitive motor neurons. eLife 8, e44423 (2019).

8. Aufderheide, K. & Whitman, M. C. Update on Congenital Cranial Dysinnervation Disorders (CCDDs). Int Ophthalmol Clin 66, 180–186 (2026).

9. Pattyn, A., Hirsch, M., Goridis, C. & Brunet, J. F. Control of hindbrain motor neuron differentiation by the homeobox gene Phox2b. Development 127, 1349–1358 (2000).

10. Jungbluth, S., Bell, E. & Lumsden, A. Specification of distinct motor neuron identities by the singular activities of individual *Hox* genes. Development 126, 2751–2758 (1999).

11. Philippidou, P. & Dasen, J. S. Hox Genes: Choreographers in Neural Development, Architects of Circuit Organization. Neuron 80, 12–34 (2013).

12. Kim, K.-T. et al. ISL1-based LIM complexes control Slit2 transcription in developing cranial motor neurons. Sci Rep 6, 36491 (2016).

13. Song, M.-R. et al. T-Box transcription factor Tbx20 regulates a genetic program for cranial motor neuron cell body migration. Development 133, 4945–4955 (2006).

14. Zeng, H. A high-resolution transcriptomic and spatial atlas of cell types in the whole mouse brain.

15. Zeisel, A. et al. Molecular Architecture of the Mouse Nervous System. Cell 174, 999–1014.e22 (2018).

16. Li, H. et al. Fly Cell Atlas: A single-nucleus transcriptomic atlas of the adult fruit fly. Science 375, eabk2432 (2022).

17. Konstantinides, N. et al. A complete temporal transcription factor series in the fly visual system. Nature 604, 316–322 (2022).

18. Seroka, A., Lai, S.-L. & Doe, C. Q. Transcriptional profiling from whole embryos to single neuroblast lineages in Drosophila. Dev Biol 489, 21–33 (2022).

19. Qiu, C. et al. A single-cell time-lapse of mouse prenatal development from gastrula to birth. Nature 626, 1084–1093 (2024).

20. Calderon, D. et al. The continuum of Drosophila embryonic development at single-cell resolution. Science 377, eabn5800 (2022).

21. Hodge, R. D. et al. Conserved cell types with divergent features in human versus mouse cortex. Nature 573, 61–68 (2019).

22. Siletti, K. et al. Transcriptomic diversity of cell types across the adult human brain. Science 382, eadd7046 (2023).

23. Farnsworth, D. R., Saunders, L. M. & Miller, A. C. A single-cell transcriptome atlas for zebrafish development. Developmental Biology 459, 100–108 (2020).

24. Farrell, J. A. et al. Single-cell reconstruction of developmental trajectories during zebrafish embryogenesis. Science 360, eaar3131 (2018).

25. Sur, A. et al. Single-cell analysis of shared signatures and transcriptional diversity during zebrafish development. Developmental Cell 58, 3028–3047.e12 (2023).

26. Saunders, L. M. et al. Embryo-scale reverse genetics at single-cell resolution. Nature 623, 782–791 (2023).

27. Alkaslasi, M. R. et al. Single nucleus RNA-sequencing defines unexpected diversity of cholinergic neuron types in the adult mouse spinal cord. Nat Commun 12, 2471 (2021).

28. Blum, J. A. et al. Single-cell transcriptomic analysis of the adult mouse spinal cord reveals molecular diversity of autonomic and skeletal motor neurons. Nat Neurosci 24, 572–583 (2021).

29. Russ, D. E. et al. A harmonized atlas of mouse spinal cord cell types and their spatial organization. Nat Commun 12, 5722 (2021).

30. Liau, E. S. et al. Single-cell transcriptomic analysis reveals diversity within mammalian spinal motor neurons. Nat Commun 14, 46 (2023).

31. Pallucchi, I. et al. Molecular blueprints for spinal circuit modules controlling locomotor speed in zebrafish. Nat Neurosci 27, 78–89 (2024).

32. D’Elia, K. P. et al. Determinants of motor neuron functional subtypes important for locomotor speed. Cell Rep 42, 113049 (2023).

33. Higashijima, S., Hotta, Y. & Okamoto, H. Visualization of Cranial Motor Neurons in Live Transgenic Zebrafish Expressing Green Fluorescent Protein Under the Control of the *Islet-1* Promoter/Enhancer. J. Neurosci. 20, 206–218 (2000).

34. Wanner, S. J. & Prince, V. E. Axon tracts guide zebrafish facial branchiomotor neuron migration through the hindbrain. Development 140, 906–915 (2013).

35. Wanner, S. J., Saeger, I., Guthrie, S. & Prince, V. E. Facial motor neuron migration advances. Curr Opin Neurobiol 23, 943–950 (2013).

36. Barsh, G. R., Isabella, A. J. & Moens, C. B. Vagus Motor Neuron Topographic Map Determined by Parallel Mechanisms of hox5 Expression and Time of Axon Initiation. Current Biology 27, 3812–3825.e3 (2017).

37. Choi, H. M. T. et al. Third-generation *in situ* hybridization chain reaction: multiplexed, quantitative, sensitive, versatile, robust. Development 145, dev165753 (2018).

38. Uemura, O. et al. Comparative functional genomics revealed conservation and diversification of three enhancers of the isl1 gene for motor and sensory neuron-specific expression. Dev Biol 278, 587–606 (2005).

39. Wang, M., Wen, H. & Brehm, P. Function of neuromuscular synapses in the zebrafish choline-acetyltransferase mutant bajan. J Neurophysiol 100, 1995–2004 (2008).

40. Ahn, D. G., Ruvinsky, I., Oates, A. C., Silver, L. M. & Ho, R. K. tbx20, a new vertebrate T-box gene expressed in the cranial motor neurons and developing cardiovascular structures in zebrafish. Mech Dev 95, 253–258 (2000).

41. Pocock, R. et al. Neuronal function of Tbx20 conserved from nematodes to vertebrates. Dev Biol 317, 671–685 (2008).

42. Szeto, D. P., Griffin, K. J. P. & Kimelman, D. HrT is required for cardiovascular development in zebrafish. Development 129, 5093–5101 (2002).

43. Duran, M. et al. A statistical framework for inferring genetic requirements from embryo-scale single-cell sequencing experiments. Preprint at 10.1101/2025.04.03.646654 (2025).

44. Barkan, E. et al. Embryo-scale single-cell chemical transcriptomics reveals dependencies between cell types and signaling pathways. Preprint at 10.1101/2025.04.03.646423 (2025).

45. Arber, S. et al. Requirement for the homeobox gene Hb9 in the consolidation of motor neuron identity. Neuron 23, 659–674 (1999).

46. Stuart, T. et al. Comprehensive Integration of Single-Cell Data. Cell 177, 1888–1902.e21 (2019).

47. Bellegarda, C., Auer, F. & Schoppik, D. Zebrafish as a model to understand extraocular motor neuron diversity. Curr Opin Neurobiol 90, 102964 (2025).

48. Greaney, M. R., Privorotskiy, A. E., D’Elia, K. P. & Schoppik, D. Extraocular motoneuron pools develop along a dorsoventral axis in zebrafish, Danio rerio. J of Comparative Neurology 525, 65–78 (2017).

49. Dowell, C. K., Hawkins, T. & Bianco, I. H. Subsets of extraocular motoneurons produce kinematically distinct saccades during hunting and exploration. Curr Biol 35, 554–573.e6 (2025).

50. Colamarino, S. A. & Tessier-Lavigne, M. The axonal chemoattractant netrin-1 is also a chemorepellent for trochlear motor axons. Cell 81, 621–629 (1995).

51. Trevarrow, B., Marks, D. L. & Kimmel, C. B. Organization of hindbrain segments in the zebrafish embryo. Neuron 4, 669–679 (1990).

52. Zannino, D. A. & Appel, B. Olig2+ Precursors Produce Abducens Motor Neurons and Oligodendrocytes in the Zebrafish Hindbrain. J Neurosci 29, 2322–2333 (2009).

53. Jurgens, J. A. et al. Gene Identification for Ocular Congenital Cranial Motor Neuron Disorders Using Human Sequencing, Zebrafish Screening, and Protein Binding Microarrays. Invest Ophthalmol Vis Sci 66, 62 (2025).

54. Asakawa, K., Abe, G. & Kawakami, K. Cellular dissection of the spinal cord motor column by BAC transgenesis and gene trapping in zebrafish. Front Neural Circuits 7, 100 (2013).

55. Bertrand, S. et al. Unexpected Novel Relational Links Uncovered by Extensive Developmental Profiling of Nuclear Receptor Expression. PLOS Genetics 3, e188 (2007).

56. Larson, J. D. et al. Expression of VE-cadherin in zebrafish embryos: A new tool to evaluate vascular development. Developmental Dynamics 231, 204–213 (2004).

57. Gershowitz, E. et al. Transcriptional profiling of extraocular motor neurons reveals sim1a as a candidate strabismus-related gene. bioRxiv 2026.04.07.717009 (2026) doi:10.64898/2026.04.07.717009.

58. Schilling, T. F. & Kimmel, C. B. Musculoskeletal patterning in the pharyngeal segments of the zebrafish embryo. Development 124, 2945–2960 (1997).

59. Diogo, R., Hinits, Y. & Hughes, S. M. Development of mandibular, hyoid and hypobranchial muscles in the zebrafish: homologies and evolution of these muscles within bony fishes and tetrapods. BMC Dev Biol 8, 24 (2008).

60. Lumsden, A. & Keynes, R. Segmental patterns of neuronal development in the chick hindbrain. Nature 337, 424–428 (1989).

61. Chandrasekhar, A., Moens, C. B., Warren, J. T., Kimmel, C. B. & Kuwada, J. Y. Development of branchiomotor neurons in zebrafish. Development 124, 2633–2644 (1997).

62. Swiatek, P. J. & Gridley, T. Perinatal lethality and defects in hindbrain development in mice homozygous for a targeted mutation of the zinc finger gene Krox20. Genes Dev. 7, 2071– 2084 (1993).

63. Schneider-Maunoury, S. et al. Disruption of Krox-20 results in alteration of rhombomeres 3 and 5 in the developing hindbrain. Cell 75, 1199–1214 (1993).

64. Torbey, P. et al. Cooperation, cis-interactions, versatility and evolutionary plasticity of multiple cis-acting elements underlie krox20 hindbrain regulation. PLOS Genetics 14, e1007581 (2018).

65. Monk, K. R. et al. A G Protein-Coupled Receptor is Essential for Schwann Cells to Initiate Myelination. Science 325, 1402–1405 (2009).

66. Sitko, A. A., Frank, M. M., Romero, G. E., Hunt, M. & Goodrich, L. V. Lateral olivocochlear neurons modulate cochlear responses to noise exposure. Proc Natl Acad Sci U S A 122, e2404558122 (2025).

67. Frank, M. M. & Goodrich, L. V. Talking back: Development of the olivocochlear efferent system. Wiley Interdiscip Rev Dev Biol 7, e324 (2018).

68. Odstrcil, I. et al. Functional and ultrastructural analysis of reafferent mechanosensation in larval zebrafish. Curr Biol 32, 176–189.e5 (2022).

69. Beiriger, A., Narayan, S., Singh, N. & Prince, V. Development and migration of the zebrafish rhombencephalic octavolateral efferent neurons. J Comp Neurol 529, 1293–1307 (2021).

70. Bricaud, O., Chaar, V., Dambly-Chaudière, C. & Ghysen, A. Early efferent innervation of the zebrafish lateral line. J Comp Neurol 434, 253–261 (2001).

71. Metcalfe, W. K., Kimmel, C. B. & Schabtach, E. Anatomy of the posterior lateral line system in young larvae of the zebrafish. J Comp Neurol 233, 377–389 (1985).

72. McArthur, K. L. & Fetcho, J. R. Key Features of Structural and Functional Organization of Zebrafish Facial Motor Neurons Are Resilient to Disruption of Neuronal Migration. Curr Biol 27, 1746–1756.e5 (2017).

73. Manuel, R., Ahemaiti, A., Tuz-Sasik, M. U. & Boije, H. Single-cell analysis of inhibitory efferent neurons of the zebrafish lateral line. PLOS ONE 21, e0346255 (2026).

74. Frank, M. M. et al. Experience-dependent flexibility in a molecularly diverse central-to-peripheral auditory feedback system. Elife 12, e83855 (2023).

75. Bingham, S., Higashijima, S., Okamoto, H. & Chandrasekhar, A. The Zebrafish trilobite gene is essential for tangential migration of branchiomotor neurons. Dev Biol 242, 149–160 (2002).

76. Wada, H. et al. Dual roles of zygotic and maternal Scribble1 in neural migration and convergent extension movements in zebrafish embryos. Development 132, 2273–2285 (2005).

77. Wada, H. & Okamoto, H. Roles of planar cell polarity pathway genes for neural migration and differentiation. Development, Growth & Differentiation 51, 233–240 (2009).

78. Mapp, O. M., Wanner, S. J., Rohrschneider, M. R. & Prince, V. E. Prickle1b mediates interpretation of migratory cues during zebrafish facial branchiomotor neuron migration. Dev Dyn 239, 1596–1608 (2010).

79. Rohrschneider, M. R., Elsen, G. E. & Prince, V. E. Zebrafish Hoxb1a regulates multiple downstream genes including prickle1b. Dev Biol 309, 358–372 (2007).

80. Mapp, O. M., Walsh, G. S., Moens, C. B., Tada, M. & Prince, V. E. Zebrafish Prickle1b mediates facial branchiomotor neuron migration via a farnesylation-dependent nuclear activity. Development 138, 2121–2132 (2011).

81. Tenney, A. P. et al. Noncoding variants alter GATA2 expression in rhombomere 4 motor neurons and cause dominant hereditary congenital facial paresis. Nat Genet 55, 1149–1163 (2023).

82. Hua, Y., Habicher, J., Carl, M., Manuel, R. & Boije, H. Novel Transgenic Zebrafish Lines to Study the CHRNA3-B4-A5 Gene Cluster. Dev Neurobiol 85, e22956 (2025).

83. Isabella, A. J. & Moens, C. B. Development and regeneration of the vagus nerve. Seminars in Cell & Developmental Biology 156, 219–227 (2024).

84. Rajendran, P. S. et al. The vagus nerve in cardiovascular physiology and pathophysiology: From evolutionary insights to clinical medicine. Seminars in Cell & Developmental Biology 156, 190–200 (2024).

85. Coverdell, T. C., Abbott, S. B. G. & Campbell, J. N. Molecular cell types as functional units of the efferent vagus nerve. Seminars in Cell & Developmental Biology 156, 210–218 (2024).

86. Kamitakahara, A., Wu, H.-H. & Levitt, P. Distinct projection targets define subpopulations of mouse brainstem vagal neurons that express the autism-associated MET receptor tyrosine kinase. J Comp Neurol 525, 3787–3808 (2017).

87. Kaneko, T., Boulanger-Weill, J., Isabella, A. J. & Moens, C. B. Position-independent functional refinement within the vagus motor topographic map. Cell Reports 43, 114740 (2024).

88. Sharma, K. et al. LIM Homeodomain Factors Lhx3 and Lhx4 Assign Subtype Identities for Motor Neurons. Cell 95, 817–828 (1998).

89. Stifani, N. Motor neurons and the generation of spinal motor neuron diversity. Front. Cell. Neurosci. 8, (2014).

90. Birkhoff, J. C., Huylebroeck, D. & Conidi, A. ZEB2, the Mowat-Wilson Syndrome Transcription Factor: Confirmations, Novel Functions, and Continuing Surprises. Genes 12, 1037 (2021).

91. Chen, T. R. et al. Generation and characterization of Tmeff2 mutant mice. Biochemical and Biophysical Research Communications 425, 189–194 (2012).

92. Rosen, Y. et al. Toward universal cell embeddings: integrating single-cell RNA-seq datasets across species with SATURN. Nat Methods 21, 1492–1500 (2024).

93. Lin, Z. et al. Evolutionary-scale prediction of atomic-level protein structure with a language model. Science 379, 1123–1130 (2023).

94. Appel, B. et al. Motoneuron fate specification revealed by patterned LIM homeobox gene expression in embryonic zebrafish. Development 121, 4117–4125 (1995).

95. Seredick, S., Hutchinson, S. A., Van Ryswyk, L., Talbot, J. C. & Eisen, J. S. Lhx3 and Lhx4 suppress Kolmer–Agduhr interneuron characteristics within zebrafish axial motoneurons. Development 141, 3900–3909 (2014).

96. Varela-Echavarría, A., Pfaff, S. L. & Guthrie, S. Differential expression of LIM homeobox genes among motor neuron subpopulations in the developing chick brain stem. Mol Cell Neurosci 8, 242–257 (1996).

97. Veerakumar, A., Yung, A. R., Liu, Y. & Krasnow, M. A. Molecularly defined circuits for cardiovascular and cardiopulmonary control. Nature 606, 739–746 (2022).

98. Isabella, A. J., Barsh, G. R., Stonick, J. A., Dubrulle, J. & Moens, C. B. Retinoic Acid Organizes the Zebrafish Vagus Motor Topographic Map via Spatiotemporal Coordination of Hgf/Met Signaling. Developmental Cell 53, 344–357.e5 (2020).

99. Henning, R. J. & Sawmiller, D. R. Vasoactive intestinal peptide: cardiovascular effects. Cardiovasc Res 49, 27–37 (2001).

100. Cuevas, J. & Adams, D. J. Vasoactive intestinal polypeptide modulation of nicotinic ACh receptor channels in rat intracardiac neurones. J Physiol 493 ( Pt 2), 503–515 (1996).

101. Braas, K. M., May, V., Harakall, S. A., Hardwick, J. C. & Parsons, R. L. Pituitary adenylate cyclase-activating polypeptide expression and modulation of neuronal excitability in guinea pig cardiac ganglia. J Neurosci 18, 9766–9779 (1998).

102. Hornung, E. et al. Neuromodulatory co-expression in cardiac vagal motor neurons of the dorsal motor nucleus of the vagus. iScience 27, 110549 (2024).

103. Stoyek, M. R., Quinn, T. A., Croll, R. P. & Smith, F. M. Zebrafish heart as a model to study the integrative autonomic control of pacemaker function. Am J Physiol Heart Circ Physiol 311, H676–688 (2016).

104. Schwerte, T., Prem, C., Mairösl, A. & Pelster, B. Development of the sympatho-vagal balance in the cardiovascular system in zebrafish (Danio rerio) characterized by power spectrum and classical signal analysis. J Exp Biol 209, 1093–1100 (2006).

105. Kuehn, E. et al. Segment number threshold determines juvenile onset of germline cluster expansion in Platynereis dumerilii. J Exp Zool B Mol Dev Evol 338, 225–240 (2022).

106. The zebrafish book: a guide for the laboratory use of zebrafish (Brachydanio rerio) - National Library of Medicine Institution. https://catalog.nlm.nih.gov/discovery/fulldisplay/alma997748323406676/01NLM_INST:01NLM_INST.

107. Siegfried, K. R. In search of determinants: gene expression during gonadal sex differentiation. J Fish Biol 76, 1879–1902 (2010).

108. Wang, J., Cao, J., Dickson, A. L. & Poss, K. D. Epicardial regeneration is guided by cardiac outflow tract and Hedgehog signalling. Nature 522, 226–230 (2015).

109. Scott, E. K. et al. Targeting neural circuitry in zebrafish using GAL4 enhancer trapping. Nat Methods 4, 323–326 (2007).

